# A multipotent cell type from term human placenta

**DOI:** 10.1101/2023.08.02.551028

**Authors:** Sangeetha Vadakke-Madathil, Esmaa Bouhamida, Bingyan J Wang, Prabhu Mathiyalagan, Parichitran Ayyamperumal, Amir Khan, Micayla Oniskey, Carlos Santos-Gallegos, Michael Hadley, Lori Croft, Fumiko Dekio, Joseph Tripodi, Vesna Najfeld, Rachel Brody, Shari Gelber, Rhoda Sperling, Hina W. Chaudhry

## Abstract

We identify a population of multipotent CDX2 cells from term human placentas with clonal expansion, migratory capacity, and immune-privileged transcriptional profiles. Isolated from 180 healthy pregnancies, these cells differentiate into cardiomyocyte and vascular lineages *in vitro* and *in vivo*. Single-cell RNA sequencing uncovers distinct cardiogenic and vasculogenic subpopulations, along with immune-modulatory and chemotactic programs, providing a blueprint for precision-guided cardiovascular cell therapy. In a NOD/SCID myocardial infarction model, CDX2 cells restore cardiac function, and clonal propagation preserves their cardiovascular differentiation potential. These findings position placental CDX2 cells as an ethically accessible, regenerative platform for targeted treatment of cardiovascular disease.

## Main

The successful use of cell therapy for cardiac repair hinges on several critical attributes: cells must have the capacity for cardiomyocyte differentiation akin to embryonic stem cells (ES) and induced pluripotent cells (iPSC), they must exhibit clonality, possess an immune-modulatory capacity for allogeneic administration, and ideally the ability to home to sites of injury. Cell therapy for cardiac purposes has been demonstrated to be safe by numerous preclinical and clinical studies ^1-4^, however; there has been a glaring lack of *bona fide* cardiac differentiation amongst cell types administered in clinical trials thus far ^5-7^. Despite a multitude of clinical cell therapy trials, which largely sought to inject stem/progenitor cells or cardiomyocytes derived from such cells into the heart, true progress in actual cardiomyocyte repopulation with a significant restoration of myocardial function has remained a distinct challenge. ES and iPS cells must be differentiated to cardiomyocytes or cardiomyocyte progenitors prior to therapeutic administration in diseased hearts, whereas we have found Caudal type homeobox-2 (CDX2) expressing cells are committed to the mesodermal fate upon isolation from human placentas. Our findings, therefore, support CDX2 cells as being naturally rooted in developmental pathways supporting organogenesis and tissue healing. Regenerative biology must rely on exploring evolutionarily preserved healing mechanisms to maintain and restore homeostasis. Gestation is one such condition that requires extensive cellular crosstalk to maintain homeostasis. The human placenta is a vital organ responsible for supporting embryonic development and facilitating the exchange of nutrients and waste between the mother and fetus. The placenta and the fetal heart are thought to undergo concurrent evolution, sharing major developmental pathways ^8-10^. Studies suggest that the placenta may regulate cardiac ventricular wall expansion once maternal blood flow is established ^11-13^. This implies that cells with regenerative ability within the placenta may be better poised to support cardiogenesis. Our focus turned to placental cells after clinical observations of patients with peripartum cardiomyopathy. This etiology of heart failure enjoys the highest recovery rate (∼50%) amongst all forms of heart failure ^14^ and we postulated that progenitor cells from the fetus or placenta might be playing a role in this recovery.

Stem or progenitor cells from the placenta are developmentally closer to embryonic stem cells but functionally different and non-tumorigenic ^15^. CDX2 is a conserved transcription factor that regulates trophoblast stem cell differentiation and is required for proper placental development ^16,17^.

In this study, we utilized 180 human term placentas from healthy patients and isolated CDX2-expressing cells from the chorionic regions. For research purposes, we utilized a human CDX2 promoter-driven lentivirus-based mCherry fluorescence reporter. Transcriptomic analyses comparing human embryonic cells, term placental CDX2 cells, and term placental non-CDX2 cells were performed to gain deeper insights into the functional differences between these cell types. We further investigated the stem/progenitor function, cellular heterogeneity at the single cell level, and cardiovascular commitment *in vitro* and *in vivo* to ascertain their potential as an innovative therapeutic for cardiovascular disease. Please refer to **Extended Data** Fig. 1 for an overview of the experimental plan.

## Results

### CDX2 cells are clonal, reside in the chorion, and harbor a cardiovascular fate signature

The homeodomain protein CDX2 is involved in embryonic development and placentation. We utilized a multi-parametric approach to detect the presence of CDX2 in human term placentas **(Fig. 1).** We examined the mRNA expression within the chorionic cells (fetal origin) of term placentas and from an earlier time point in first-trimester chorionic villus samples (CVS). CDX2 mRNA was detected in chorionic regions of term placentas and CVS samples (**Fig. 1a** and raw data in **Extended Data** Fig. 2a, b, primer sequences in **Supplementary Table 1).**

**Fig. 1:**
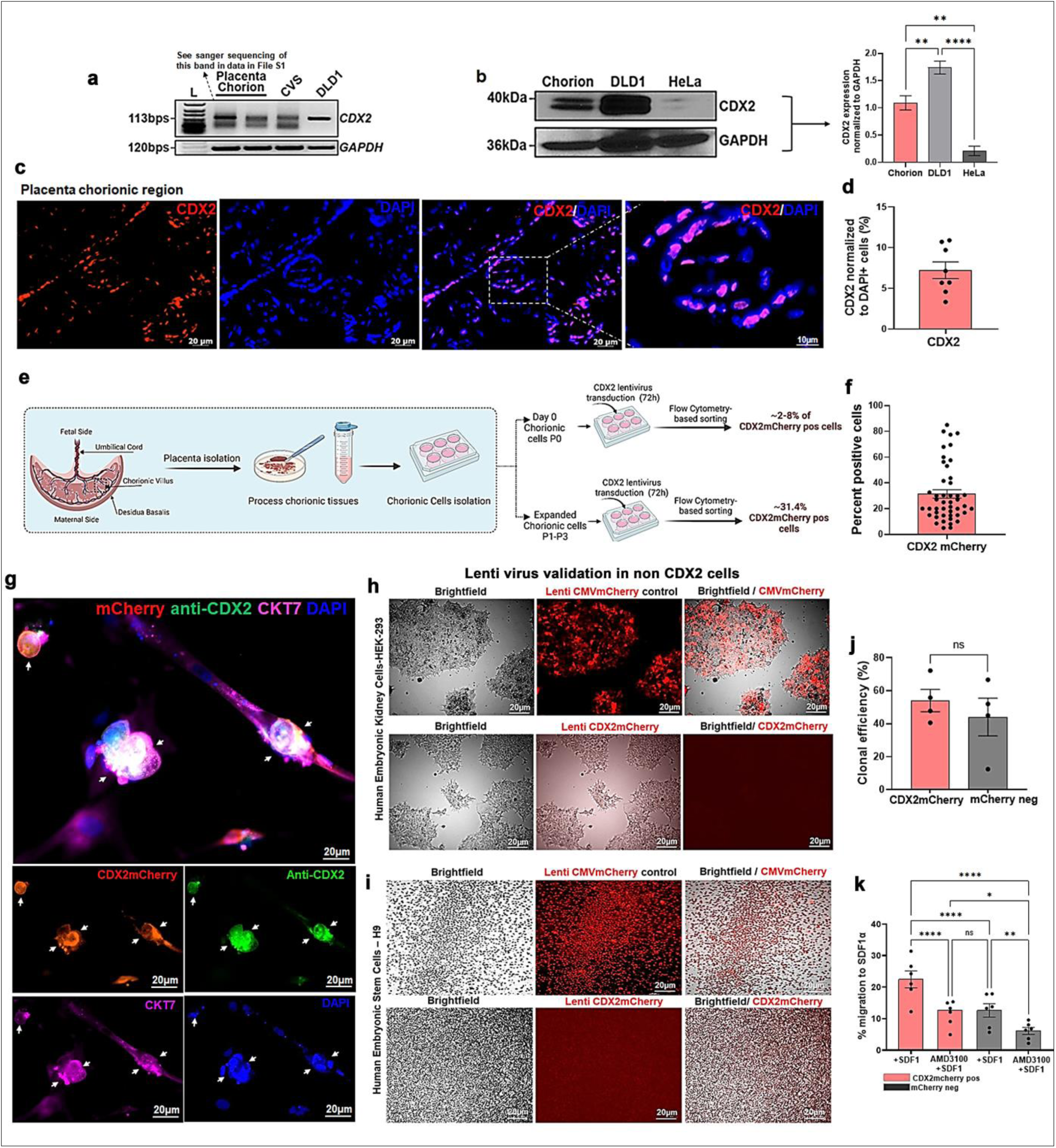
CDX2 expression, lentivirus-based isolation, validation and progenitor function. **a)** CDX2 mRNA expression in the first-trimester chorionic villous sample and term chorion from the healthy placenta. Two different placental samples showing CDX2 mRNA expression with bands showing splice variants. Human adenocarcinoma cell line DLD1 shows a higher expression of endogenous *CDX2 and* hence used as a positive control. *GAPDH* was used as an internal control. L=DNA ladder. **b)** Western blot analyses of human term chorionic sample showing CDX2 protein expression with bands showing splice variants. DLD1 was used as a positive control, while HeLa cells were used as a control with low endogenous CDX2 expression. Protein expression normalized to GAPDH shown to the right. Data are represented as mean ± SEM, **p<0.01, ****p<0.0001, n=2. **c)** Immunofluorescence images showing CDX2 expression in the term cytotrophoblast cells in the term chorion. **d)** Quantification of tissue sections showing the percentage of CDX2 expressing cells in the FFPE tissue. Data are represented as mean ± SEM, n=4 independent samples, dot plot represents technical replicates (2 ROI each). **e)** Schematic showing chorionic cell isolation, plating and lenti virus transduction to select CDX2 expressing cells. Transduction on P0 (passage 0 = after isolation and partial confluence), yielded ∼2-8% of CDX2 cells. **f)** Culturing the chorionic cells to P1-P3 and further transduction yielded ∼31.4% ± 3.2 of CDX2 cells. Data are represented as mean ± SEM, n=46 independent placenta samples. **g)** Specificity of the lentivirus-based selection of mCherry cells was validated using co-localization of mCherry fluorescence with CDX2 (Green) antibody (CDX-88clone) and trophoblast marker Cytokeratin 7 (CKT7-pink). **h, i)** Immunofluorescence images showing two non-CDX2 expressing cell types (HEK293 and H9ES), transduced using CDX2mCherry lentivirus alongside a CMV-driven constitutively active mCherry lentivirus (control). Each row of top panels (with columns for brightfield alone, mCherry alone, and merged brightfield/mCherry) clearly shows CMV mCherry fluorescence in identical experimental conditions. Bottom row of panels reveal no endogenous CDX2 expression validating the specificity of our CDX2 lentiviral reporter system. **j)** Bar graph showing the clonal efficiency of sorted CDX2mCherry and mCherry negative cells on day 14 (54.1 ± 6.78% in CDX2 cells versus 44 ± 11.45% in mCherry negative cells). Data are presented as mean ± SEM (n=4 independent samples). **k)** Bar graph showing the enhanced *in vitro* migration of CDX2mChery cells in response to SDF1α (vs. mCherry negative cells). Blocking CXCR4 with AMD3100 significantly reduced the migration, showing that the migration is partly mediated by SDF1-CXCR4 axis. Data are presented as mean ± SEM. n=3 independent samples with 3 technical replicates per sample is shown,*p<0.05, **p<0.01.

Sanger sequencing and analysis of the output FASTA sequences confirmed sequences specific to human CDX2 mRNA (**Source Data Fig.1**). Immunoblot analyses detected CDX2 protein in the term chorion (∼40kDa) with the conventional doublet pattern representing two isoforms from alternative splicing^18,19^ (**Fig. 1b a**nd raw data in **Extended Data** Fig. 2c, d). HeLa cells showed a lower endogenous expression of CDX2 while human adenocarcinoma cell line DLD1 served as a positive control. Immunohistochemistry revealed that CDX2 protein was enriched in the fetal chorionic region of the human placenta and not in the fetal amnion **Extended Data** Fig. 2e, f). CDX2 was detected in the villous region **(Fig. 1c**, merged enlarged region within the boxed ROI, and **Fig. 1d**), particularly within the cytotrophoblast layer of the chorion. De-identified human colon tumor tissue sections and human DLD1 cells served as a positive control and displayed a higher CDX2 expression (**Extended Data** Fig. 2g, h).

The isolation of viable CDX2 cells from human placental tissue is challenging due to the lack of known surface markers for enrichment. To overcome this, we developed a lentiviral reporter system driven by the human CDX2 promoter to enable selective labeling and isolation of CDX2-expressing cells based on mCherry fluorescence (**Extended Data** Fig. 3a–c; see Methods). Among three promoter constructs tested, a 612 bp fragment upstream of the CDX2 transcriptional start site (designated Promoter 1) drove robust mCherry expression in DLD1 cells and was selected for further use (**Extended Data** Fig. 3d–g). Transduction efficiency was evaluated across a range of multiplicities of infection (MOI 20–100) in DLD1 cells (**Extended Data** Fig. 3h–q). An MOI of 50 was chosen as optimal, balancing transduction efficiency and minimal cytotoxicity. Using this system, chorionic cells isolated from term placentas were transduced with the CDX2-promoter: mCherry lentivirus, allowing for the prospective identification and enrichment of CDX2mCherry cells based on the downstream fluorescence. Following isolation and initial plating, adherent chorionic cells from term placentas were transduced with the selected CDX2 promoter–driven mCherry lentiviral construct (hereafter referred to as CDX2mCherry cells; **Fig. 1e**). At baseline, only 2–8% of freshly isolated chorionic cells exhibited detectable CDX2 expression. After 1–3 *in vitro* passages, transduction of the expanded adherent population and subsequent cell sorting via flow cytometry enabled a mean frequency of ∼31.4% ± 3.2 **(Fig. 1f),** to be isolated for downstream analyses.

Immunofluorescence analysis confirmed the specificity of the CDX2 selection by demonstrating co-localization of mCherry signal (red) with nuclear CDX2 (green) in transduced chorionic cells (**Fig. 1g and Extended Data** Fig. 2i). These CDX2mCherry cells also stained positive for the pan-trophoblast marker cytokeratin 7 (pink), indicating a trophoblast identity and confirming that the mCherry lentivirus truly marks endogenous CDX2-expressing cells (**Fig. 1g and Extended Data** Fig. 2j). To further validate transcriptional specificity, we compared CDX2 promoter–driven expression to a constitutively active CMV promoter in non-CDX2-expressing HEK293 cells and H9 embryonic stem (H9ESCs) cells. CMV mCherry lentivirus yielded robust fluorescence, confirming successful transduction (**Fig. 1h, i, top panels**). In contrast, no mCherry signal was detected in HEK293 and H9 ESCs transduced with the CDX2mCherry construct (**Fig. 1h, i**, bottom panels), indicating lack of CDX2 expression in those cells and the promoter specificity.

To evaluate the self-renewal potential of CDX2mCherry cells, we assessed their clonal proliferative capacity over a 14-day period following single-cell isolation. CDX2mCherry cells exhibited robust clonal expansion with an efficiency of 54.1% ± 6.78% **(Fig. 1j)**. Interestingly, mCherry negative chorionic cells also demonstrated clonal proliferation, albeit at a slightly lower efficiency (44% ± 11.45%). This is not surprising considering that the presence of multiple progenitor populations, aside from CDX2, have been reported within the placenta^20-22^. Similarly, transcriptional profiling of non-CDX2 (mCherry negative) cells revealed an elevated expression of hematopoietic lineage markers, including the pan-leukocyte marker PTPRC (CD45), the myeloid progenitor marker CD33, and CD1b, suggesting a predominantly hematopoietic commitment **(Extended Data** Fig. 3r–t). Given the incomplete efficiency of lentiviral transduction, it is also likely that a fraction of CDX2mCherry cells were not labeled, as reflected in the positive control DLD1 cell line **(Extended Data** Fig. 3u). To assess the progenitor functional response of CDX2mCherry cells to chemotactic signaling, we performed a transwell migration assay using stromal-derived factor 1α (SDF1α). CDX2mCherry cells showed significantly enhanced migration toward SDF1α (22.4% ± 2.68%) compared to mCherry negative cells (12.62% ± 2.11%; **Fig. 1k**). Pre-treatment with AMD3100, a CXCR4 antagonist, substantially reduced migration, indicating that the chemotactic response is at least partly mediated via the SDF1–CXCR4 axis. Collectively, these results support a stem/progenitor-like phenotype of placental CDX2 cells, characterized by clonal self-renewal and chemotactic responsiveness.

CDX2 is a key regulator of trophectoderm lineage specification and is essential for trophoblast-mediated placentation. In contrast, human embryonic stem cells (hESCs), which are derived from the inner cell mass, show marked downregulation of CDX2 expression **(Fig. 1i).** To delineate lineage-specific transcriptional programs further, we performed bulk RNA sequencing of CDX2mCherry placental cells, mCherry negative placental cells, and H9 hESCs **(Source Data Fig. 2a** and **Source Data Fig. 2b**). Differential expression analyses revealed 1,065 genes significantly upregulated and 154 genes downregulated in CDX2mCherry cells relative to hESCs. When compared to the mCherry negative placental population, CDX2mCherry cells exhibited 46 upregulated and 529 downregulated genes **(Supplementary Table 2).** These findings suggest a transcriptional distinctiveness of CDX2-expressing placental cells, distinguishing them from both pluripotent embryonic stem cells and hematopoietic-like placental populations.

**Fig. 2:**
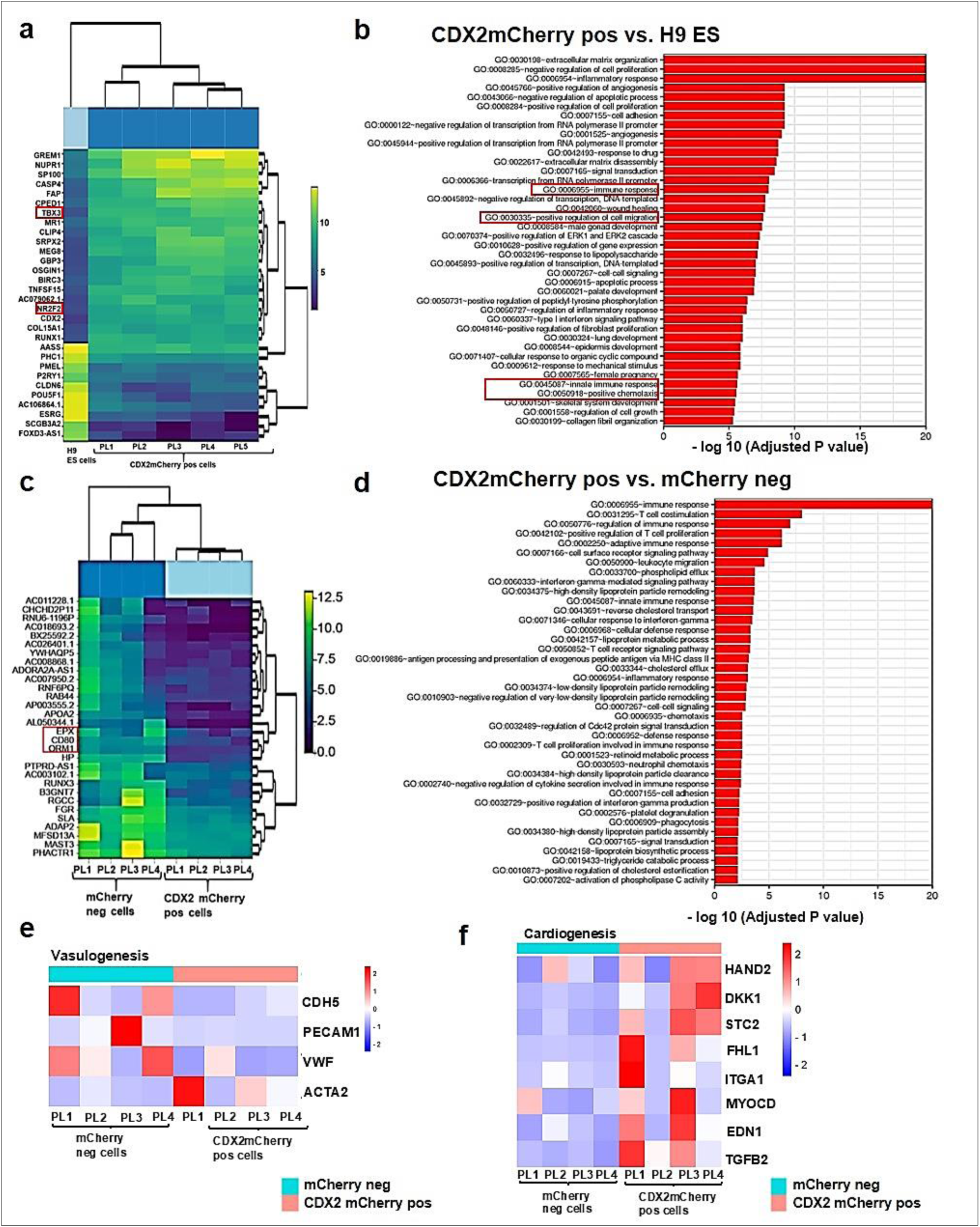
Bulk RNAseq analyses showing human placental CDX2 cells are uniquely poised for cardiovascular commitment. **a)** Heatmap showing the expression profile of the top 30 differentially expressed genes in CDX2 cells (5 independent samples labeled as PL1-PL5) versus hESCs (H9 ES ctrl from 3 pooled samples-extreme left). Genes were ranked by their adjusted p-values, and log₂-transformed expression values were plotted. The yellow-green color code indicates upregulated genes, and the dark blue-gray color code indicates downregulated genes across the samples. CDX2mCherry cells showed upregulated cardiac mesoderm commitment genes *(NR2F2* and *TBX3,* highlighted in the red box) while downregulating pluripotent markers (*CLDN6*, *POU5F1,* and *ESRG).* **b)** Bar plot showing the pathways enriched in CDX2mCherry cells vs. H9ES cells corresponding to functions related to angiogenesis, immune modulation, migration, and chemotaxis. **c)** Heatmap with the top 30 differentially expressed genes showing the upregulated function of inflammatory response and immune genes in mCherry negative cells (n=4 independent samples in each group). **d)** Bar plot showing the pathways analyses demonstrating upregulation of functional pathways including T cell proliferation, MHC class II antigen processing, and presentation, IFN gamma mediated signaling, and cholesterol biosynthesis in mCherry negative cells. **e, f)** Heatmap showing the expression of vasculogenic genes *(PECAM1, CDH5, vWF, and ACAT2) and cardiogenic genes (HAND2, DKK1, STC2, FHL1, ITGA1, MYOCD, EDN1,* and *TGFB2)* in CDX2 vs mCherry negative cells.

To examine lineage-specific transcriptional differences, we generated a biclustering heatmap of the top 30 differentially expressed genes (DEGs) between CDX2mCherry placental cells and H9ESCs **(Fig. 2a).** Functional enrichment analysis of these DEGs revealed significant associations with pathways involved in tissue organization, cell differentiation, angiogenesis, cell survival, and cardiac development. Notably, *TBX3* and *NR2F2*, genes implicated in mesodermal specification and early cardiogenesis, were selectively upregulated in CDX2mCherry cells but downregulated in hESCs, suggesting a transcriptional bias of CDX2 cells toward an extraembryonic mesodermal and cardiovascular lineage program.

In contrast, genes associated with pluripotency—including *POU5F1, FOXD3-AS1, CLDN6*, and *PHC1*—were significantly downregulated in CDX2mCherry cells relative to H9 hESCs **(Fig. 2a)**, lowering the risk of tumorigenicity often linked to undifferentiated hESCs and iPSCs ^23-27^. Gene ontology (GO) analysis further indicated that CDX2mCherry cells are enriched for pathways involved in immune modulation and angiogenesis **(Fig. 2b).** Moreover, gene sets related to cell migration and chemotaxis were significantly enriched in CDX2mCherry cells compared to H9 hESCs, consistent with their enhanced migratory capacity observed in transwell assays **(Fig. 1k**, also refer to **Source Data Fig. 2a** supplement document). Analysis of the transcriptome of mCherry negative placental cells revealed upregulation of genes involved in immune activation and inflammatory responses **(Fig. 2c, d**, also refer to **Source Data Fig. 2b).** These included *RAB44*, a Ras-family gene implicated in immune cell signaling, as well as *EPX* and *CD80*, a costimulatory molecule essential for T and B cell activation. Acute-phase proteins such as *ORM1, HP,* and *APOA2*; the latter involved in HDL metabolism and linked to atherosclerotic processes were also elevated in this population.

Gene expression analysis revealed a differential expression of vasculogenesis-associated transcripts in the CDX2mCherry population, including *PECAM1 (CD31), CDH5 (VE-cadherin), VWF, and ACTA2* **(Fig. 2e)**, specifying an angiogenic/vascular gene program. Further pathway enrichment analysis uncovered upregulation of genes involved in endothelial-to-mesenchymal transition (EndMT) and valvulogenesis, including *SNAI2, KLF4, YAP1, VCAN (Versican), EGFR, CD44, TGFB2*, and *BMP4* in CDX2mCherry cells compared to the mCherry negative fraction **(Extended Data** Fig. 4a, b). Notably, *NOTCH2* and *GSK3B* expression levels were significantly elevated in the CDX2mCherry population (**Extended Data** Fig. 4a, b; *p < 0.05), further supporting a shift towards mesenchymal and valvular lineage trajectories. Strikingly, other distinct clusters of genes implicated in mesodermal specification and early cardiac lineage commitment—including *HAND2, DKK1, STC2, FHL1, ITGA, MYOCD, EDN1,* and *TGFB2*—were preferentially expressed in CDX2mCherry cells **(Fig. 2f)**. These transcripts are known regulators of cardiac development and structural remodeling in both fetal and adult heart tissues. Gene sets associated with sarcomere organization and cardiac muscle function, including *FHL1, STC2, DKK1, TGFB2,* and *EDN1,* were significantly upregulated relative to mCherry negative cells from the same placental samples **(Fig. 2f).** Collectively, these data demonstrate that CDX2mCherry cells from the term chorion are transcriptionally distinct from undifferentiated pluripotent stem cells and are farther along the mesodermal/cardiac commitment pathway and are enriched for a committed cardiovascular gene signature.

### CDX2mCherry cells differentiate into endothelial cells and smooth muscle cells *in vitro* while they display neo-angiogenesis *in vivo*

We noted that CDX2mCherry cells can develop endothelial vessel-like structures (27.3% ±5.6) and express endothelial markers PECAM1 (CD31) and von Willebrand factor (vWF) when co-cultured with murine neonatal cardiomyocytes **(Fig. 3a-c).** Under feeder-free conditions, these cells exhibited uniform expression of the smooth muscle marker SM22α (95.5% ± 1.59%), indicative of mesodermal lineage commitment **(Fig. 3d–e).** We examined angiogenic functionality wherein CDX2mCherry cells were subjected to a Matrigel-based tube formation assay. Within 3 hours of seeding, CDX2 cells extended tubular projections, and by 6 hours, formed endothelial-like networks (**Fig. 3f, j**; refer to **Movie S1** in loop mode). mCherry negative cells produced less organized structures under identical conditions (**Fig. 3g, k**). HUVECs, included as a positive control, formed robust and interconnected capillary-like networks at both time points (**Fig. 3h–l**; refer to **Movie S2** in loop mode). Unfractionated chorionic cells did not form tubular structures **(Fig. 3i, m).**

**Fig. 3:**
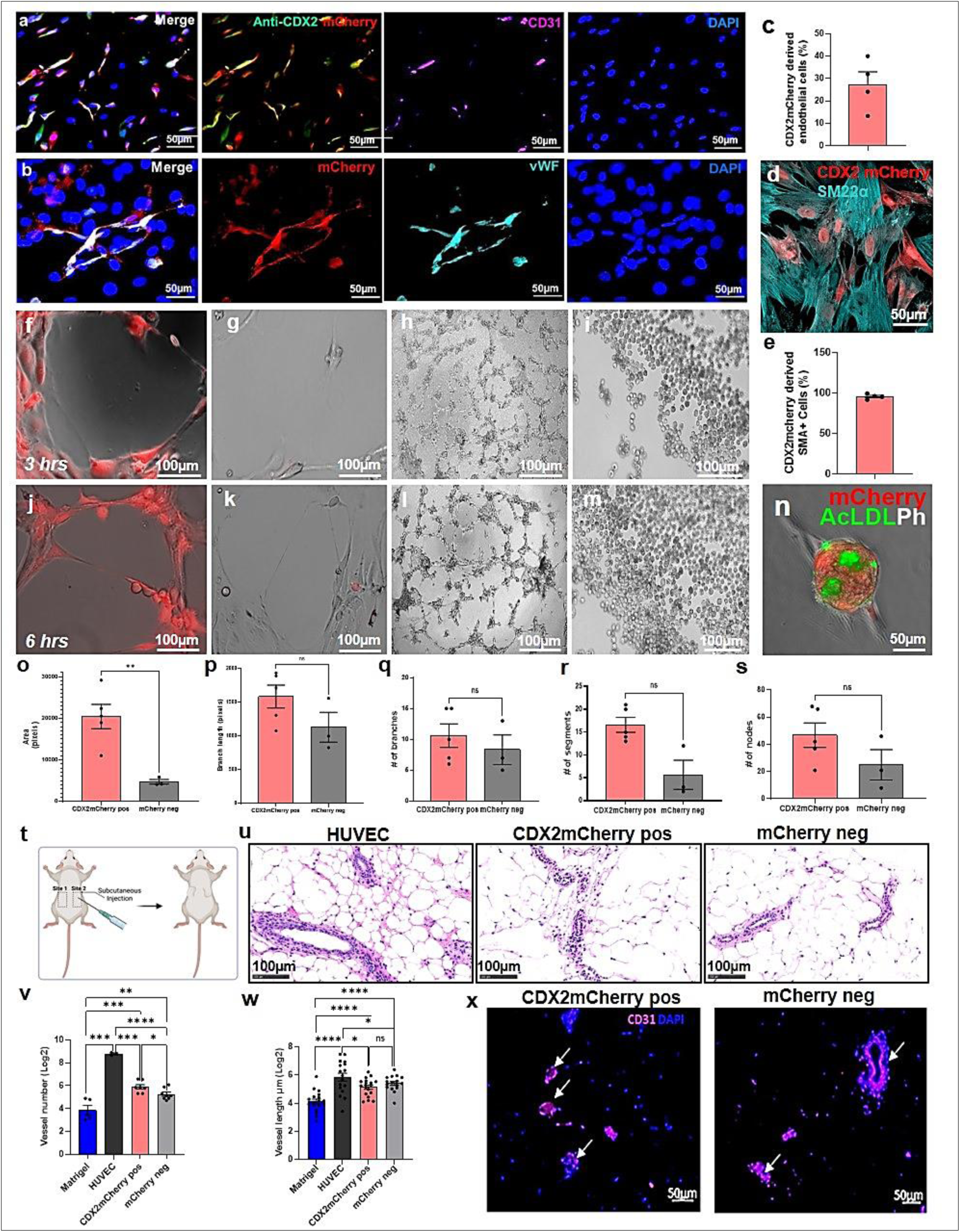
CDX2mCherry cells are angiogenic and display endothelial tube formation *in vitro* and *in vivo*. **a**) Immunofluorescence images show a subset of CDX2mCherry cells (red) express vascular lineage marker CD31 (pink) and give rise to blood vessel-like assembly.**b)** Immunofluorescence staining of endothelial marker von Willebrand Factor (vWF-green) upon co-culture with neonatal cardiomyocytes. **c)** Bar plots showing quantification of CDX2mCherry negativederived endothelial cells. Data represented as mean ± SEM (n=4). **d)** CDX2mCherry cells also expressed smooth muscle marker (SM22α) in feeder-free culture using IMDM+10%FBS. **e)** Quantification of immunofluorescence data represented as mean ± SEM (n=4). **f-m)** Endothelial tube formation on Matrigel in 3hrs (all top panels) and 6hrs (all bottom panels). **f)** CDX2mCherry cells forming endothelial network as imaged at 3hrs and **j)** at 6hrs. *Refer to movie S1 in loop mode.* **g, k)** mCherry negative cells exhibited less organized networks compared to those formed by CDX2mCherry cells on Matrigel. **h, l)** Human umbilical vein endothelial cells (HUVEC) served as a positive control; also *refer to Movie S2 in loop mode*. **i, m)** Total chorionic cells did not show endothelial tube formation at both time points. **n)** Ac-LDL (Alexa-488) uptake by primed (M200 medium +LSGS+ 10ng/mL VEGF) CDX2mCherry cells suggesting the formation of a functionally active endothelium. **o-s)** Quantification of the endothelial network using ImageJ software (angiogenesis analyzer plugin) showed a higher endothelial area (loop) and branching segments in CDX2 cells compared to mCherry negative cells. Data represented as mean ± SEM, *p<0.05 (CDX2mCherry n=5 and mCherry negative n=3 samples). **t**) Overview of the experimental plan of the Matrigel plug assay for *in vivo* angiogenesis assessment (n=3 per group). **u)** Representative microscopic images of hematoxylin and eosin (H&E) (20× magnification) of HUVEC, CDX2mCherry cells, and mCherry negative cells. **v)** Bar graph showing the vessel numbers, and **w)** the vessel length in HUVEC cells, CDX2mCherry cells, and negative cells. CDX2 cells showed a significant increase in vessel numbers, *p≤0.05. Data represented as mean ± SEM (dots in the bar chart indicate areas (ROIs) quantified in each group. Vessel numbers-Matrigel=5, HUVEC=3, CDX2mCherry=6, and mCherry negative=8. Vessel length-Matrigel=18, HUVEC=17, CDX2mCherry=18, and mCherry negative=16. **x)** Representative immunofluorescence images showing CD31 (purple) and DAPI (blue) staining in both CDX2 and mCherry negative cells derived vessels.

To assess functional endothelial activity, CDX2mCherry cells were primed in endothelial growth medium supplemented with 10 ng/mL vascular endothelial growth factor (VEGF) and incubated with acetylated low-density lipoprotein (Ac-LDL alexa 488) for 4 hours. Robust Ac-LDL uptake was observed, confirming receptor-mediated endocytosis and endothelial functionality **(Fig. 3n)**. Quantitative analysis of network formation^28^ revealed that CDX2mCherry cells formed significantly more mesh structures compared to mCherry negative cells (*p ≤ 0.05; **Fig. 3o**), with a trend toward increased total branch length, number of segments, and nodes (**Fig. 3p–s**). To evaluate *in vivo* angiogenic potential, undifferentiated CDX2mCherry cells were embedded in Matrigel and subcutaneously implanted into NOD/SCID mice **(Fig. 3t)** alongside controls including HUVEC+ Matrigel, mCherry negative cells+ Matrigel and Matrigel alone (no cells). After 14 days, histological analysis of explanted plugs revealed vessel-like structures originating from both CDX2mCherry and mCherry negative groups, though less extensive than in HUVEC-containing plugs, which exhibited much bigger and an organized vasculature **(Fig. 3u).** CDX2mCherry cells contributed to elongated vessel-like structures within and surrounding the Matrigel plugs. Quantification showed a significantly greater number of vessels in CDX2mCherry plugs compared to mCherry negative controls (63 ± 8.7 vs. 40 ± 5.3; *p ≤ 0.05; **Fig. 3v**), while vessel length did not differ significantly (**Fig. 3w**). Immunostaining confirmed human-specific CD31 expression in vessels derived from both CDX2mCherry and mCherry negative cells **(Fig. 3x)**. These findings demonstrate that CDX2mCherry placental cells possess functional endothelial characteristics *in vitro* and contribute to neoangiogenesis *in vivo*, supporting their potential to differentiate toward endothelial and vascular lineages.

### CDX2mCherry cells robustly differentiate into spontaneously beating cardiomyocytes *in vitro* and significantly improve cardiac function after myocardial infarction

To evaluate the cardiomyogenic potential of CDX2mCherry cells, we utilized a neonatal murine cardiomyocyte feeder-based system previously established in murine studies as a supportive niche for cardiomyocyte differentiation without exogenous inducers^22,29,30^.

CDX2mCherry cells isolated from healthy term human chorions were plated onto spontaneously beating neonatal cardiomyocyte feeders. Time-lapse live-cell imaging revealed the formation of beating, organoid-like clusters by day 4–5 in culture (**Fig. 4a**; refer to **Movie S3** in loop mode). By two weeks, these clusters exhibited spreading and rod-shaped morphologies while maintaining spontaneous contractility (**Fig. 4b**; refer to **Movie S4** in loop mode). After four weeks of co-culture, immunostaining confirmed expression of human-specific sarcomeric α-actinin and cardiac troponin T (cTnT, cross-reactive to both human and murine cells), with clear sarcomeric striations, indicating robust cardiomyocyte differentiation **(Fig. 4c–e’)**. In contrast, mCherry negative cells cultured under identical conditions displayed limited cardiomyogenic potential, with significantly lower differentiation efficiency **(5.09% ± 1.38%**; **Extended Data** Fig. 5a–e). Quantitative analysis showed that ∼71.11% ± 12.15 of input CDX2mCherry cells adopted a cardiomyocyte phenotype **(Fig. 4f),** highlighting their strong propensity for cardiac lineage commitment. To rigorously exclude the possibility of cell fusion with murine feeder cardiomyocytes, we performed fluorescence in situ hybridization (FISH) using species and sex chromosome-specific nuclear probes. Human CDX2-derived cardiomyocytes were identified as diploid, bearing either XX chromosomes labeled with green (491–516 nm) or orange (525–551 nm) human-specific probes. Murine feeder nuclei were distinctly labeled using aqua XX probes (418–467 nm) (**Fig. 4g).** FISH analysis confirmed that CDX2-derived cardiomyocytes were diploid and human in origin, exhibiting only a single set of sex chromosomes, thereby eliminating the possibility of cell fusion.

**Fig. 4:**
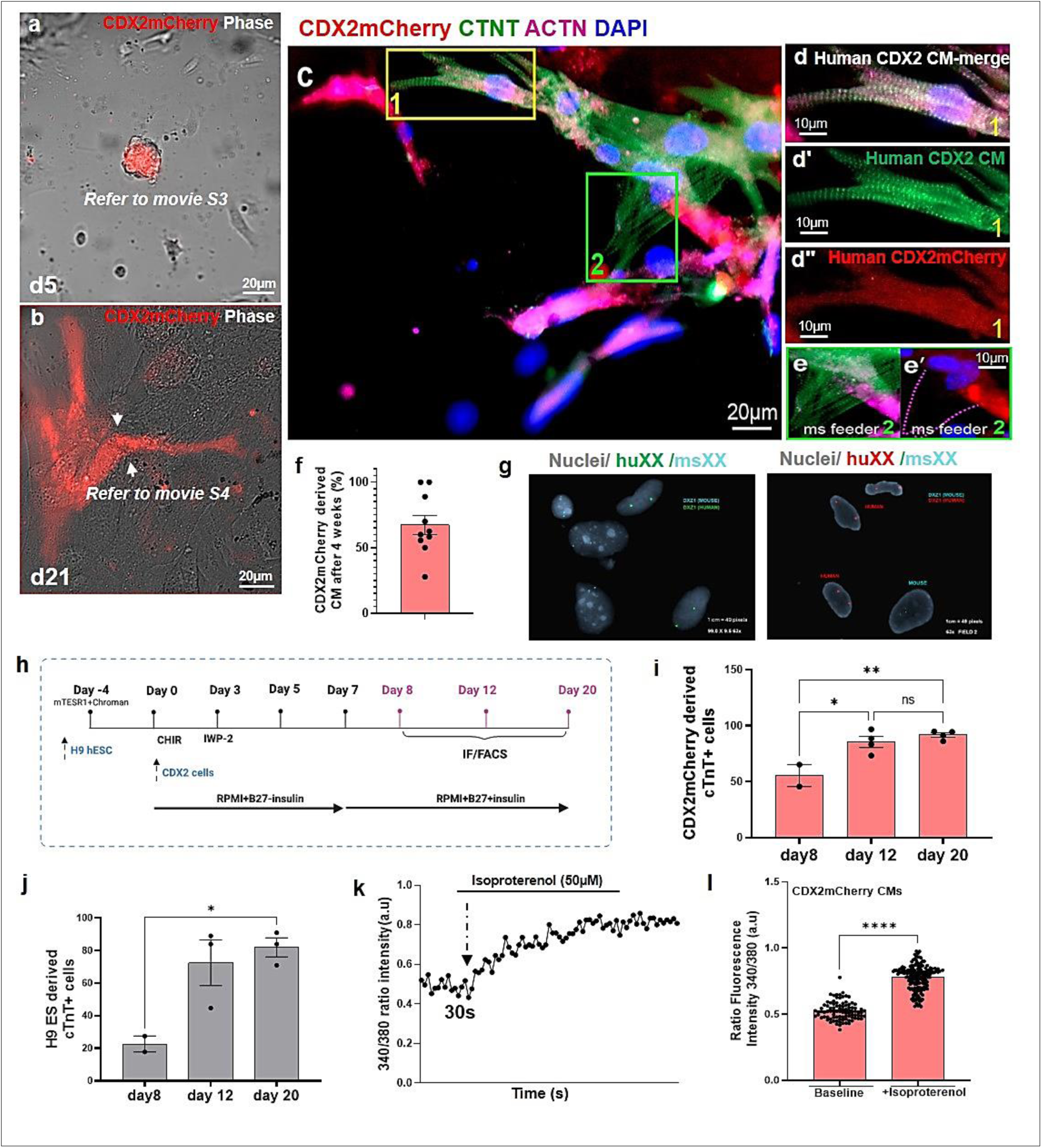
CDX2mCherry cells differentiate into spontaneously beating cardiomyocytes. **a)** Snapshot of a beating CDX2mcherry cell cluster (day 5) in co-culture with murine neonatal cardiomyocytes feeder (non-mCherry-gray cells in the background), refer to *movie S3 in loop mode.* **b)** Snapshot of beating-adhered and rod shaped CDX2 derived cardiomyocytes 3 weeks in culture. Refer to *movie S3 in loop mode.* **c)** CDX2-derived cardiomyocytes (mCherry negativered) stained positive for human cardiac sarcomeric actinin (pink-Alexa 647) and (cTnT-green Alexa 488). The specificity of the human actinin antibody was detected by a lack of staining on murine feeder myocytes in the background, while cTnT (green) detects both mouse and human cardiomyocytes. Yellow box 1 area is enlarged to the right in **d-d’** showing the striated sarcomeres. **d”)** mCherry negative fluorescence. Green box 2 and the enlarged panels **e, e**’ point to the endogenous murine feeder myocyte that expresses only cTnT (green). Nuclei are counterstained using DAPI (blue). **f)** Bar plot showing the quantification of CDX2mCherry derived cardiomyocytes. Approximately 71.11% ± 12.15 of input CDX2mCherry cells differentiated into cardiomyocytes. Data are represented as mean ± SEM (n=5 independent samples, with dot plots reflecting 2 ROIs analyzed per sample). **g)** Fluorescence Insitu hybridization (FISH) images showing the presence of only one set of sex chromosomes in CDX2-derived cardiomyocyte nuclei, emphasizing that CDX2 derived CMs did not exhibit cell fusion with murine feeder myocytes (n=3). Human cells are diploid and were detected using two different human-specific probes (XX green-dUTP 491nm-516nm) and XX (Orange-dUTP 525nm-551nm), whereas XX aqua probes (Aqu-dUTP 418nm-467nm) identified murine feeder cells. **h)** Overview of feeder-free cardiac differentiation of CDX2mCherry cells using temporal modulation of Wnt signaling. H9ES cells were used as controls under identical culture conditions. **i, j)** Cardiac-specific marker cTnT to quantify differentiation on day 8, day 12 and day 20. CDX2mCherry cells showed an efficient and early differentiation, especially on day 8, in comparison to H9ES cells at the same time points. Data represented as mean ± SEM, *p<0.05, **p<0.01. H9ES n=2 day 8 and n=3 for day 12 and day 20, CDX2mCherry n=2 day 8 and n=4 for day 12 and day 20. **k, l)** Calcium measurement in CDX2-derived cardiomyocytes after day 20 of feeder free differentiation using Fura2-AM ratiometric measurements. Panel **k** shows the intensity ratio in baseline (30s) and the increase in the ratio in response to Isoproterenol from a representative sample. Panel **l** represents the data of 3 samples pooled and analyzed in triplicate showing significant increase in calcium spike (baseline 93 readings and Isoproterenol 165 readings in the dot plot) after Isoproterenol treatment in CDX2-derived CMS p<0.0001, mean ±SEM.

To assess whether CDX2mCherry cells could be directed toward cardiomyocyte differentiation in a feeder-free system, we employed a modified GiWi protocol involving the temporal modulation of Wnt signaling using small-molecule inhibitors, as previously described^31^. H9ESCs were included as a positive control under identical culture conditions. Based on prior findings from our co-culture system, we focused exclusively on the CDX2mCherry population for induction. As outlined in the schematic **(Fig. 4h)**, CDX2mCherry cells were seeded on day 0, and differentiation was assessed at days 8, 12, and 20 post-induction. At each time point, we evaluated cardiomyocyte differentiation by immunostaining and flow cytometry quantification for cardiac troponin T (cTnT). Differentiation efficiency increased progressively over time. The percentage of cTnT^+^ cardiomyocytes derived from input CDX2mCherry cells was ∼55.45% on day 8, 85.5% on day 12, and 91.7% by day 20. **(Fig. 4i)**. Representative flow cytometry plots illustrate this progressive increase in cTnT^+^ cells **(Extended Data** Fig. 5f), which was accompanied by a reduction in mCherry fluorescence, consistent with lineage commitment and differentiation. Interestingly, we also observed inter-sample variability in mCherry retention following differentiation. In some samples, a subset of cTnT^+^ cardiomyocytes continued to express mCherry fluorescence even at day 20 **(Extended Data** Fig. 5g, h) suggesting inherent biological heterogeneity among CDX2 cells from different placental donors.

Under identical culture conditions, H9ESCs exhibited cardiomyocyte differentiation, with cTnT^+^ cells comprising ∼22.65%, 72.5%, and 82% on days 8, 12, and 20, respectively **(Fig. 4j).** Notably, CDX2mCherry cells outperformed H9 hESCs at early stages of differentiation, with significantly higher frequencies of cTnT+ cardiomyocytes by day 8: 55.45% cTnT^+^ in CDX2 cells versus 22.65% in H9ESCs, (*p < 0.05; **Fig. 4i–j**). These findings suggest a more rapid and efficient cardiogenic commitment in the CDX2 population. To assess functional maturation, we examined intracellular calcium handling in CDX2-derived cardiomyocytes on day 20 using Fura-2 AM calcium imaging. Ratiometric traces demonstrated clear basal calcium transients and robust responses to isoproterenol, a β-adrenergic agonist **(Fig. 4k, l)**, indicating that CDX2-derived cardiomyocytes possess functional calcium flux response. Together, these results demonstrate that CDX2mCherry cells from human term placenta can be efficiently directed toward the cardiac lineages using two different culture conditions. Compared to hESCs, they show a faster response in differentiation, reinforcing their potential as a readily accessible, non-embryonic source of cardiomyocyte progenitors.

To directly evaluate the therapeutic potential of human placental CDX2mCherry cells in cardiac repair, we employed a mouse model of myocardial infarction (MI) using NOD/SCID mice (9–9.5 weeks old, both sexes; **Fig. 5a**). MI was induced by permanent ligation of the left anterior descending (LAD) coronary artery, using previously established protocols ^22,30^. Immediately following LAD ligation, mice received intramyocardial injections of CDX2mCherry cells (∼100,000 cells in 5 µL per injection site) at three equidistant locations around the peri-infarct region (n = 7). A separate cohort received mCherry negative cells under identical conditions (n = 7), and a vehicle control group received phosphate-buffered saline (PBS) injections (n = 6). Cardiac magnetic resonance imaging (MRI) was performed the day after surgery and injection, to establish a post-MI baseline, and repeated one month later to assess structural and functional outcomes **(Fig. 5a).**

**Fig. 5:**
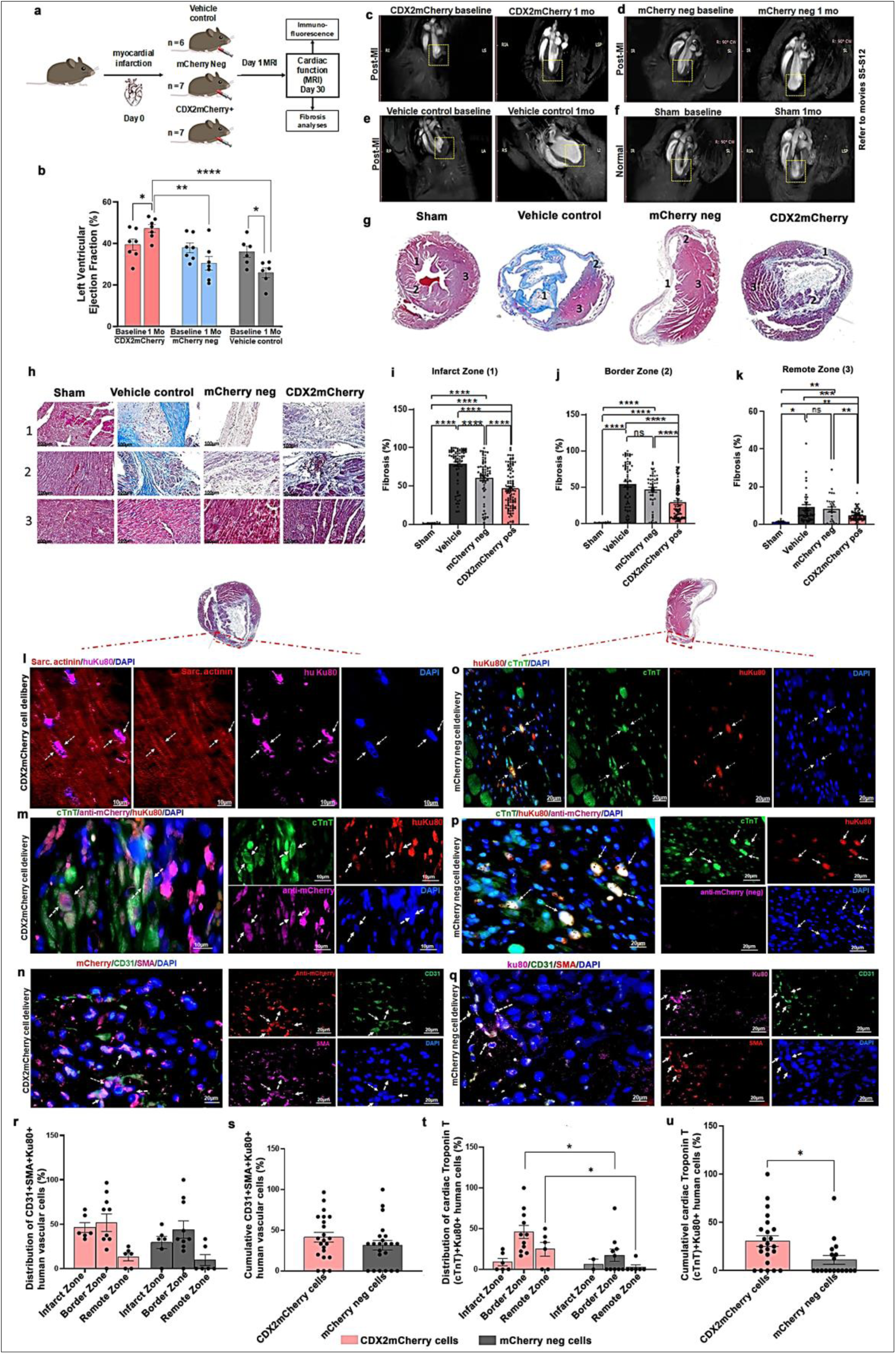
CDX2 cell delivery significantly improved cardiac function after myocardial infarction. **a)** Experimental design for myocardial infarction, cell delivery and analyses in NOD/SCID mice. **b)** Cardiac MRI analyses showing a significant increase in left ventricular ejection fraction (EF) in CDX2 cell-injected mice versus the control mice at 1-mo time points compared with the baseline MI timeline (CDX2 group, n = 7; mCherry negative group, n = 7, Vehicle control PBS n=6). Data are represented as mean ± SEM. *p≤0.05 (CDX2 baseline MI vs. 1 mo post-MI and Veh ctrl baseline MI vs. I mo post MI), **p≤0.01 (CDX2 at 1 mo vs. mCherry negative at 1 mo post-MI) and ****p≤0.0001 (CDX2 1mo post-MI vs. PBS group 1 mo post-MI). **c)** Long axis view of cardiac MRI showing reduced ventricular wall dilation (boxed areas in all images) upon CDX2 cell delivery in 1-mo compared to **d)** mCherry negative cell group, **e)** the vehicle control group, and **f)** Sham mice that do not undergo MI show a normal ventricular wall at both time points, as shown. Please see movies S5-S12 in loop mode. **g)** Trichrome analyses to visualize MIs and to evaluate fibrosis percentage. Representative myocardium from sham (no MI), vehicle control (MI+PBS injection), MI+ mCherry negative cell injection and MI+CDX2 cell injection groups (3 slides from each of 3 mice/group examined). Fibrotic areas are stained blue (collagen), and viable tissue is stained red (muscle). **h)** Images of Infarct zone, IZ (1), Border zone, BZ (2), and Remote zone, RZ (3) are shown with panels, **i-k)** presenting quantified fibrosis percentages. Data show a significant reduction in fibrosis in CDX2 cell-treated groups compared to both mCherry negative cell injected and PBS injected mice post MI. Data are represented as mean ± SEM (n=3) with each dot representing the cumulative quantified ROIs per area (Sham = 10 ROIs; CDX2mCherry and mCherry negative = 30-70 ROIs per zone, *p<0.05, **p<0.01, **p<0.001 and ***p<0.0001). **l)** Engraftment and differentiation of CDX2 cells into cardiomyocytes in the post-MI heart. A representative image from the boxed region in the border zone shows CDX2-derived cardiomyocytes (CDX2 CMs, white arrows) exhibiting striated sarcomeric structures with co-expression of sarcomeric α-actinin (red) and human Ku80 (huKu80, pink). **m)** Alternate field of CDX2 derived cardiomyocytes (white arrows) expressing cTnT (green), anti-mCherry (pink) and huKu80 (red). **n)** CDX2mCherry cells contributed to vascular differentiation in vivo. Representative image shows CDX2mCherry cells associated with vascular structures co-expressing CD31 and smooth muscle actin (SMA), indicating endothelial and smooth muscle lineage differentiation. **o, p)** Limited cardiac differentiation of mCherry negative cells in the injured myocardium. mCherry negative cells are identified by human Ku80 (red), and cardiomyocytes are immunostained for cTnT (green). Panel p represent an additional field confirming a lack of mCherry expression in mCherry negative cells. **q)** mCherry negative cells contributed to vascular differentiation (ku80-pink, SMA-red and CD31-green) cells. **r)** Bar plot showing quantification of CDX2 and mCherry negative derived vascular cells in infarct, border and remote zones. Each dot in the graph represents a region of interest (ROI) analyzed from the infarct zone (IZ, n = 6), border zone (BZ, n = 10), and remote zone (RZ, n = 6) in both groups. **s)** Quantification of overall vascular cell contribution by CDX2 and mCherry negative cells. **t)** Bar plot quantifying CDX2mCherry and mCherry negative cell-derived cardiomyocytes in the infarct (IZ), border (BZ), and remote (RZ) zones. CDX2mCherry cells show significantly higher cardiogenic potential post-MI compared to mCherry negative cells. ROI counts: CDX2 group – IZ = 6, BZ = 11, RZ = 6; mCherry negative group – IZ = 6, BZ = 10, RZ = 6. *p ≤ 0.05. **u)** Quantification of overall cardiomyocyte formation by CDX2 and mCherry negative cells in the post MI heart. Data are presented as mean ± SEM.

Clinical cardiologists with expertise in cardiac MRI, who were blinded to the treatment groups, assessed cardiac function. MRI analysis revealed a significant improvement in contractile function in the CDX2mCherry cell–treated group, as measured by increased left ventricular ejection fraction (LVEF), compared to both vehicle (PBS) controls and the mCherry negative cell group **(Fig. 5b)**. In the CDX2 group, LVEF improved from 39.89% ± 2.6 at baseline (day 1 post-MI) to 47.26% ± 1.8 at one month (*p ≤ 0.05). In contrast, vehicle-treated mice and mCherry negative cell-treated mice showed a decline in LVEF (Vehicle treated 25.9% ± 2.3, and mCherry negative cell-treated mice 30.4% ± 3.2 of EF, **p<0.01). Comparisons at the one-month time point demonstrated superior functional recovery in the CDX2 group relative to both controls (****p ≤ 0.0001 vs PBS; **p ≤ 0.01 vs mCherry negative). Representative long-axis MRI frames **(Fig. 5c–f)**, refer to **Movies S5–S12** in loop mode), illustrate the preservation of ventricular wall thickness, and reduced chamber dilation in CDX2mCherry treated hearts compared to controls. Individual LVEF values for each mouse are presented in **Supplementary Table 3.** It is important to note that NOD/SCID mice are highly susceptible to MI-induced mortality, limiting the severity of infarction that can be induced ^32^. Post-MI LVEF in this model cannot be lowered more than ∼35–40%, as more extensive damage typically results in procedural mortality prior to cell administration. To assess structural remodeling, we analyzed fibrosis in transverse heart sections using Masson’s Trichrome staining one-month post-MI. Fibrosis was quantified in the infarct zone, border zone, and remote zone in the injured myocardium from each of the three animals per group all sectioned just below mid-ventricle **(Fig. 5g–k)**. CDX2mCherry cell–treated hearts exhibited a significant reduction in fibrotic area compared to both mCherry negative, and vehicle controls: Infarct zone: CDX2 group: 46.3% ± 2.5, vs. mCherry negative: 60.4% ± 3.1, and PBS: 78.6% ± 2.7 (****p < 0.0001), Border zone: CDX2: 28.4% ± 2.4, vs. mCherry negative: 46.4% ± 3.3, and PBS: 54.0% ± 3.8 (****p < 0.0001), and Remote zone: CDX2: 4.6% ± 0.3, vs. mCherry negative: 7.8% ± 1.3, and PBS: 8.9% ± 1.4 (**p < 0.01). As expected, sham-operated hearts had minimal fibrosis (1.3% ± 0.2%). Notably, CDX2-treated sections displayed areas of intact healthy myocardium along both endocardial and epicardial borders of the infarct, a feature absent in control groups. This histological evidence, together with the functional data, supports a regenerative role for human placental CDX2 cells in promoting myocardial repair post-infarction.

To investigate whether functional cardiac improvement following CDX2 cell delivery was associated with lineage-specific differentiation, we performed immunohistochemical analysis of NOD/SCID mouse hearts. Human-specific Ku80 and mCherry were used as markers to identify engrafted human cells. In the border zone of the infarct, (highlighted dotted boxed regions) CDX2mCherry cells differentiated into cardiomyocytes with sarcomeres visualized by co-expression of sarcomeric actinin (actinin red, white arrows, **Fig. 5l).** A 3D orthogonal view showing the nuclei and Ku80 fluorescence within the actinin+ cardiomyocyte is shown in **Figure S5b** and cardiac troponin T (cTnT, green, **Fig. 5m**). These cells also expressed Ku80 and mCherry (pink), confirming their human origin (white arrows). Additional supporting images are provided in **Extended Data** Fig. 6a, b. Furthermore, CDX2mCherry cells demonstrated vascular lineage differentiation, co-expressing smooth muscle actin (SMA) and CD31, an endothelial marker (**Fig. 5n**, anti-mCherry red, CD31 green, SMA pink). While mCherry negative cells (identified using anti-Ku80) were present in the myocardium, they exhibited minimal cardiomyogenic differentiation (**Fig. 5o, p**) and limited contribution to new cardiomyocytes (**Extended Data** Fig. 6c). To ensure specificity, anti-mCherry antibody staining was applied to both experimental and control tissues. As expected, mCherry negative cell–treated hearts showed no detectable mCherry signal **(Fig. 5p)**. However, evidence of vascular differentiation was observed in these cells (**Fig. 5q**), albeit to a lesser extent than in the CDX2mcherry group. These findings collectively demonstrate that human CDX2 placental cells engraft within the infarcted heart and contribute to both cardiac and vascular cell lineages, supporting their role in direct tissue regeneration and repair.

Quantification of CDX2-derived vascular cells in the infarct, border, and distal zones revealed a trend toward higher expression of CD31 and SMA in CDX2mCherry cells compared to mCherry negative cells. However, this difference was not statistically significant. CDX2mCherry cells accounted for 46.25% ± 5.56, 51.74% ± 9.83, and 13.06% ± 4.3 of CD31+ SMA+ cells in the infarct, border, and distal zones, respectively. Overall contribution was 41.32% ± 5.9; while mCherry negative cells contributed to 29.4% ± 6.9, 44.03% ± 9.81, and 9.72% ± 6.2 of CD31+SMA+ cells in the infarct, border, and distal zones, respectively, yielding an overall vascular contribution of 31.6% ± 5.84 **(Fig. 5r, s)** one month after cell delivery in the post-MI heart.

CDX2mCherry cells, however, exhibited significantly enhanced cardiogenic capacity. Following delivery into the infarcted heart, CDX2mCherry (mCherry+/Ku80+) cells co-expressed cTnT and accounted for 8.83% ± 4.54, 45.83% ± 7.9, and 24.72% ± 8.5 of cTnT+ cardiomyocytes in the infarct, border, and distal zones, respectively. mCherry neg cells, by comparison, demonstrated minimal cardiomyogenic differentiation (6.25% ± 6.25, 17.22% ± 7.5, and 2.7% ± 2.7 in the corresponding zones; **Fig. 5t, u**, *p<0.05-CDX2 groups in border and remote zones vs. mCherry negative groups). CDX2mCherry-derived cardiomyocytes displayed hallmarks of maturity, including organized sarcomeric architecture, and localized predominantly within the injury and border zones, regions critical for functional recovery as noted by MRI. Flow cytometric analysis of non-cardiac organs in NOD/SCID recipients showed no detectable human cell engraftment **(Extended Data** Fig. 6d), supporting the safety and tissue specificity of CDX2mCherry cell delivery. Together, these findings position human placental CDX2mCherry cells as multipotent progenitors capable of both vasculogenic and cardiogenic differentiation *in vivo*. The mCherry negative fraction appeared restricted to a vascular fate with negligible cardiomyogenic output, consistent with transcriptomic and *in vitro* differentiation profiles **(Fig. 2a, e, f).** Functionally, CDX2 cell therapy led to reduced fibrosis, enhanced neovascularization and cardiomyocyte regeneration, and significantly improved cardiac function, underscoring its potential for regenerative myocardial repair.

### Single cell RNA sequencing reveals unique cellular subsets within CDX2mCherry cells further supporting a role for cardiovascular commitment

While culturing CDX2mCherry cells, we noticed the overall retention of mCherry fluorescence through passages **(Fig. 6a-c)** with a significant increase in cell yield after each passage until approximately 1 million cells were generated by passage 6 based on input cell number 50,000 from only 5-gram chorionic tissue (noting that average chorionic plate weighs ∼150 g). To evaluate the scalability of CDX2mCherry cell expansion, we conducted a proof-of-concept experiment using the BioOne1250 2L single-use benchtop bioreactor system (Distek BioOne, Distek Inc., NJ, USA). Cells were cultured on Corning Synthemax II microcarriers for 10 days. CDX2mCherry cells demonstrated efficient attachment and robust proliferation within the bioreactor environment, with cell density increasing from 7.5 × 10^5 cells/mL to 5.0 × 10^6 cells/mL **(Fig. 6d),** representing an approximately 7-fold expansion. Fluorescence microscopy confirmed increased cell attachment and aggregation on microcarriers over time **(Fig. 6e, f).** Even in the absence of bioreactor optimization, these findings establish the feasibility of scalable CDX2mCherry cell expansion to clinically relevant levels.

**Fig. 6:**
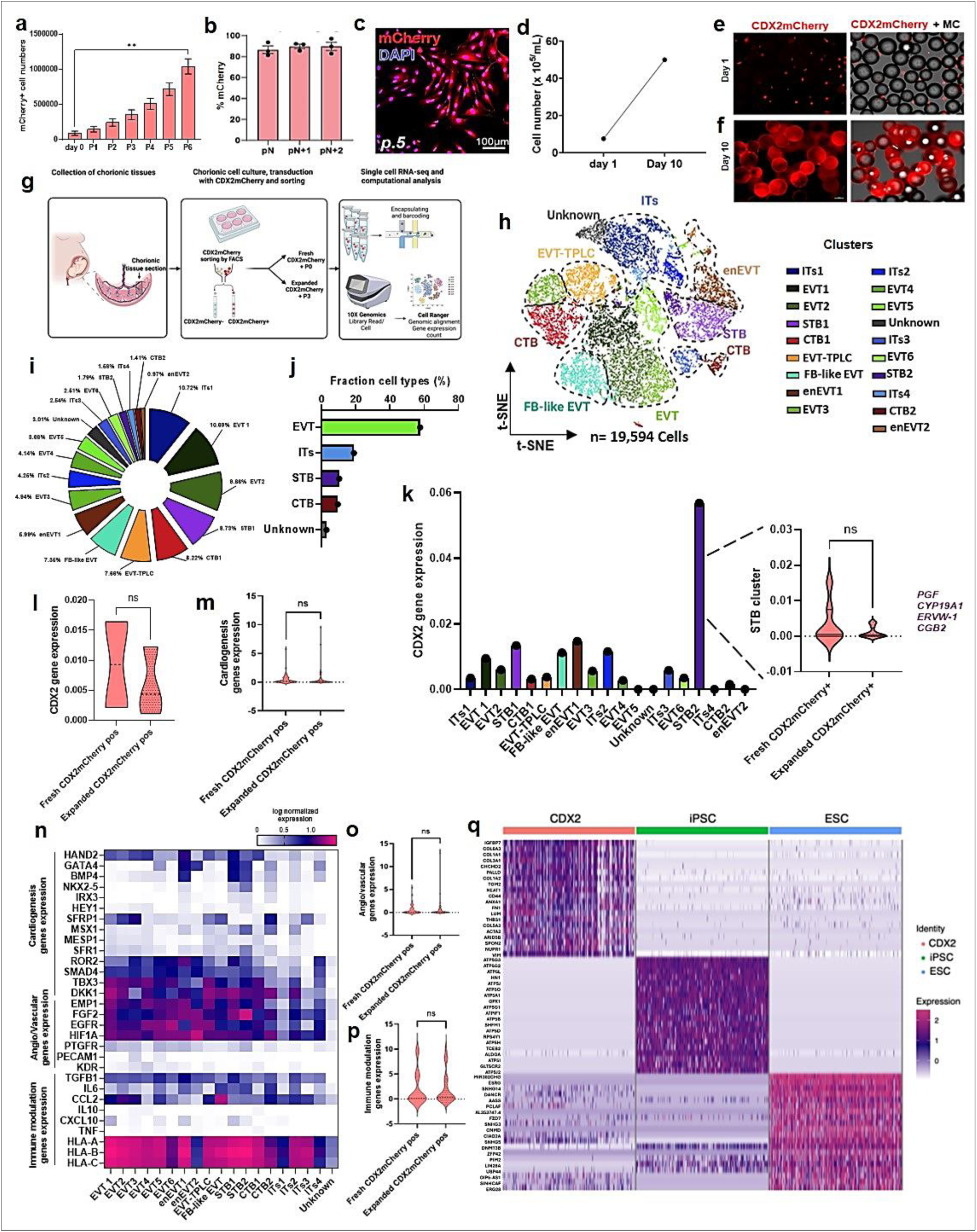
Ex vivo expansion and scRNA-seq reveals unique cellular subclusters with cardiovascular commitment in placental CDX2 cells. **a)** Quantification of mCherry cells after expansion showed a significant increase, i.e., up to 1 million cells by passage 6, compared to day 0 of isolation. Data are represented as mean ± SEM (n=3, **p<0.01). b) Retention of mCherry expression over various passages as analyzed by flow cytometry (n=3), and **c)** representative fluorescent microscopy image at passage 5 (p5). **d)** Line plot showing the cell yield of CDX2mCherry cells on day 1 versus day 10 in the bioreactor culture. **e)** Representative fluorescence microscopy images showing a remarkable difference in cell attachment and proliferation of CDX2mCherry cells (mCherry) on synthemax II microcarriers at the beginning (1 day) and **f)** at the end (day 10) of bioreactor-based expansion. Data are representative of pooled CDX2mCherry cells isolated and expanded from 3 different placenta samples. **g)** Schematic representation of the workflow for isolating and analyzing human placental CDX2mCherry cells using scRNA-seq. **h)** T-distributed Stochastic Neighbor Embedding (t-SNE) plot of human placental CDX2mCherry cells (fresh n=2 and expanded n=3). Each dot represents a single cell, with different colors indicating distinct cell types (19 clusters). Villous cytotrophoblasts (CTB), syncytiotrophoblasts (STB), extravillous trophoblasts (EVT), Endovascular extravillous trophoblast (enEVT), EVT-trophoblast progenitor-like cells (EVT-TPLC), Fibroblasts like trophoblasts (FB-like EVT), Intermediate trophoblasts (ITs) and Unknown cells. **i)** Donut chart showing the percentages of CDX2 subclusters. **j)** Bar plot showing the fraction of major cell types. **k)** Bar graph depicting CDX2 expression levels across the subclusters. STB2 cluster, exhibit higher CDX2 expression and is present in both fresh and expanded populations. **l)** Violin plot showing no significant downregulation of CDX2 gene expression. **m)** Violin plots illustrate overall cardiogenic gene expression levels in fresh versus expanded CDX2mCherry cells. **n)** Heatmap showing the expression levels of cardiogenic, angiogenic/vasculogenic, and immune-modulatory genes across various clusters. **o)** Violin plot showing the expression levels of angiovascular genes, and **p)** immune modulatory genes in fresh and expanded CDX2mCherry cells, refer to the gene lists provided in supplementary Data File S5 for panel m, o and p. **q)** Row-normalized heat map of the top 20 marker genes for each cell type - CDX2⁺ cells, iPSCs, and ESCs—identified via single-cell RNA sequencing. Publicly available iPSC and ESC datasets were integrated for comparison. CDX2⁺ cells exhibited unique upregulation of extracellular matrix-related genes and genes involved in cardiac-related functional pathways, including TGM2, ACTA2, SPON2, and AXIN2, among others. (Refer to supplementary data file S6 for the complete list of differentially expressed genes supporting the distinction of CDX2⁺ cells from iPSC and ESC populations).

To assess the translational potential of CDX2mCherry cells for therapeutic applications, we investigated their cellular heterogeneity and cardiovascular gene expression programs, particularly in the context of *ex vivo* expansion—an essential step for clinical scalability. We performed single-cell RNA sequencing (scRNA-seq) using the 10x Genomics platform on freshly isolated (n = 2) and ex vivo-expanded CDX2mCherry cells (passage 3, n = 3) **(Fig. 6g).** Unsupervised clustering via t-distributed stochastic neighbor embedding (t-SNE) analysis revealed a broad and unbiased distribution of 19,594 single cells pooled from both fresh and expanded samples **(Fig. 6h)**. Cell annotation was performed using cell marker identification based on *classical trophoblast gene markers* ^33-36^ and the upregulated gene expression profiles from our dataset. Based on transcriptional similarities, we identified 19 distinct cellular subpopulations (**Extended Data** Fig. 7a-c; also refer to **Source Data Fig. 6a**). Gene expression profiling enabled the classification of specific trophoblast subtypes within the CDX2mCherry population **(Fig. 6i, j)**. Notably, extravillous trophoblasts (EVTs), marked by *PSG1* and *DIO2*, constituted the majority (57.77%) of cells. Subsets of EVTs included endovascular extravillous trophoblasts (enEVTs; *PRDM6*) and fibroblast-like EVTs (FB-like EVTs; *COL1A1, CCN2*). We also identified extravillous trophoblast progenitor-like cells (EVT-TPLCs; *MKI67, TOP2A*), villous cytotrophoblasts (CTBs; *KRT7, EFEMP1*) accounting for 9.63% of the population, and syncytiotrophoblasts (STBs; *PGF, CGB2*) representing 10.51%. Intermediate trophoblasts— cells with mixed or transitional expression profiles between EVT, CTB, and STB lineages—comprised approximately 19% of the total population, while a small fraction (3%) remained unclassified due to ambiguous marker expression.

CDX2 expression was maintained in expanded CDX2mCherry cells, with no significant downregulation observed when compared to freshly isolated samples. CDX2 expression was predominantly higher in the STB2 cluster **(Fig. 6k, l).** Transcriptomic profiling revealed that CDX2mCherry cells expressed a panel of genes associated with cardiac mesoderm development, including *HAND2, GATA4, BMP4, ROR2, SMAD4, TBX3, DKK1, MSX1, MESP1*, *SFRP1, NKX2-5, KDR, HEY1, IRX3,* and *TBX6.* scRNA-seq also corroborated bulk RNA-seq findings, confirming enrichment of gene programs associated with epithelial-to-mesenchymal transition (EMT/EndMT) and valvular development in the CDX2mCherry population **(Extended Data** Fig. 7d, e). Importantly, there were no significant differences in cardiac gene expression profiles between fresh and expanded CDX2mCherry samples (**Fig. 6m**, also refer to **Source Data Fig. 6b**), with the exception of *SFRP1*, which was significantly upregulated in the expanded population (**Fig. 6n** and **Extended Data** Fig. 8a-d **\*\***p<0.01). *SFRP1* is a known antagonist of the Wnt/β-catenin signaling pathway^37^, and its elevated expression in expanded cells was consistent with bulk RNA-seq results. Expression of angiogenesis- and vasculogenesis-related genes remained stable between fresh and expanded cells **(Fig. 6o**, also refer to Data S5 supplement). However, expression of cardiac mesoderm genes was heterogeneous across cellular subpopulations. Notably, *NKX2-5, IRX3, MSX1, GATA4, BMP4, HAND2, DKK1*, and *SFRP1* were highly expressed within STB clusters, particularly STB2 **(Fig. 6n)**. These genes are critical regulators of early cardiogenesis, playing essential roles in mesodermal specification and cardiomyocyte differentiation. In contrast, EVT clusters—including EVT1–EVT6, enEVT2, EVT-TPLC, and FB-like EVT—exhibited low to negligible expression of *NKX2-5,* with enEVT1 showing the lowest levels. Nevertheless, other cardiac-associated genes such as *HAND2, ROR2, SMAD4, TBX3, SFRP1,* and *DKK1* were variably expressed across EVT clusters. Notably, EVT3 subcluster displayed minimal *DKK1* expression, suggesting a bias toward a more angiogenic commitment. CTB clusters expressed a smaller subset of cardiac mesoderm genes but still retained expression of *HAND2, GATA4, BMP4, MSX1, SMAD4, ROR2, and SFRP1.* Furthermore, genes involved in angiogenesis and vasculogenesis—including *PECAM1, EGFR, PTGFR, KDR, HIF1A,* and *EMP1*—were highly expressed across EVT clusters, while *FGF2* was notably enriched in the STB2 cluster **(Fig. 6n),** aligning with the observed cardiac vs. vascular functional bias of these subpopulations.

Within the CDX2mCherry cell population, the STB2 cluster emerged as highly enriched for genes critical to cardiac mesoderm development, including *FGF2*, underscoring its potential role in cardiomyocyte differentiation. Notably, STB2 exhibited the highest levels of CDX2 expression among all subclusters and co-expressed *NKX2-5,* a definitive marker of cardiac progenitor identity. The STB2 cluster also exhibited higher levels of *MYC* and *KLF4*, transcription factors linked to cell proliferation, reprogramming, differentiation, and cardiogenesis, highlighting the unique molecular identity of this subset. In contrast, EVT populations were predominantly associated with angiogenic and vasculogenic gene signatures. Specifically, EVT3 lacked expression of key cardiogenic genes such as *NKX2-5, IRX3,* and *GATA4*, and exhibited only minimal levels of *DKK1* and *BMP4*, suggesting a transcriptional program skewed toward vascular lineage commitment. We also identified a distinct EVT subpopulation, designated EVT-TPLC, characterized by a highly proliferative gene signature. This cluster showed strong expression of canonical cell cycle and mitotic regulators, including *MKI67, KIF23, TOP2A, ANLN, AURKA, AURKB, TEAD4, CCNA2, CDK1*, and *PCNA* **(Extended Data** Fig. 9a, b), indicating a transcriptional state consistent with active proliferation.

To investigate the immunomodulatory potential of placental cells, we profiled the expression of immune-associated genes, including *TGFB1, CXCL10, CCL2, IL6, IL10, TNF,* and *HLA-A,* - *HLA-B,* and *HLA-C*. Among these, *CCL2* was markedly upregulated in the FB-like EVT population, implicating a role in chemotaxis and immune regulation **(Fig. 6n)**. Expression levels of these genes did not significantly differ between freshly isolated and expanded CDX2mCherry cells **(Fig. 6p**, also refer to **Source Data Fig.5a**). At single-cell resolution, several surface markers were found to be enriched in CDX2mCherry cells, including *SERPINE1, HMMR, CD9, and AQP1 (***Extended Data** Fig. 9c–f). Notably, *AQP1*, which facilitates ion transport and cell migration, exhibited elevated expression in specific STB subsets. *HMMR*, a regulator of cell motility, extracellular matrix remodeling, and proliferation, was predominantly expressed in STB clusters, particularly STB2 and the progenitor cluster EVT-TPLC, suggesting potential utility as a marker for this distinct population **(Extended Data** Fig. 9d). Additionally, beyond *CXCL12* and *CXCR4*, our scRNA-seq analysis identified other candidate homing-associated genes i.e., *VCAM1, ICAM1, ACKR3, SIRP1, HEG1 and ITGA4* **(Extended Data** Fig. 9g), which may inform strategies to optimize targeted homing of CDX2mCherry cell subsets in future applications.

We examined the differentiation potential and lineage relationships of CDX2mCherry cells, using pseudotime trajectory analysis to reconstruct cell-state transitions across integrated scRNA-seq datasets. Our dataset of CDX2mCherry cells was analyzed alongside previously published single-cell profiles^38-40^ of human iPSCs and their differentiated progeny, including cardiomyocytes (CMs) and endothelial cells (ECs) (**Extended Data** Fig. 10). Pseudotime mapping revealed that CDX2mcherry cells align more closely with iPSC-derived CMs and ECs, suggesting a developmental trajectory where CDX2mCherry cells may serve as precursors to cardiomyocyte and endothelial lineages. Subsets of CDX2mCherry cells mapped to distinct points along this differentiation continuum, reflecting transcriptional heterogeneity within the population. Cells located earlier in pseudotime were transcriptionally similar to iPSCs (orange box), while those positioned further along displayed profiles resembling D5 ECs (blue box) or D30 CMs (green box), consistent with a multipotent, lineage-primed state.

Notably, CDX2mCherry cells exhibited minimal transcript level overlap with undifferentiated iPSCs and ESCs, indicating that they represent a molecularly distinct population other than the pluripotent stem cells (**Fig. 6q** and refer to **Source Data Fig. 6c**). Differential gene expression analysis identified a CDX2 specific signature enriched for extracellular matrix components *COL6A3, COL1A1, COL3A1, PALLD, FN1, CD44, LUM* and genes implicated in cardiac homeostasis, including *TGM2, SPON2, ANXA1,* and *THBS1^41-44^* In contrast, pluripotency-associated transcripts such as *ZFP42, PIM2, LIN28A*, and *USP44^45-48^* were selectively downregulated in CDX2mCherry cells compared to iPSCs and ESCs. Taken together, these findings suggest that human placenta-derived CDX2 cells represent a transcriptionally distinct, multipotent population with developmental trajectories biased toward cardiovascular lineages, positioning them as promising candidates for cardiovascular regenerative applications.

## Discussion

In this study, we show that CDX2-expressing cells isolated from term human placentas exhibit fundamental properties of stem/progenitor cells, including the ability to migrate in response to the chemokine SDF1α and to differentiate into cardiovascular lineages. CDX2, a transcription factor critical for early embryonic development, is expressed in the trophectoderm— the precursor to the placenta. Notably, placental and cardiac development occur in parallel during embryogenesis, forming a placenta-heart axis in which the placenta influences fetal cardiomyocyte proliferation^8-13^. Our findings support this inter-organ connection by demonstrating that placenta-derived CDX2 cells are uniquely primed for cardiovascular differentiation. These cells now emerge as promising candidates for driving cardiogenesis and may serve as unique cellular targets for regenerative cardiac therapies.

For decades, a variety of cell sources have been explored for cardiac regeneration, with many demonstrating safety in clinical trials. Yet, identifying an ideal, readily accessible cell type—capable of differentiating into cardiovascular lineages outside of ESCs and iPSCs— remains a major challenge. Prenatal placental cells, such as placental mesenchymal stem cells (PMSCs) and amniotic epithelial cells (AECs), offer promising anti-inflammatory, angiogenic, and paracrine effects, but they have not been shown to generate *de novo* cardiomyocytes^49,50^. In contrast, endogenous trophoblast progenitors within the placenta have been largely overlooked for their potential to differentiate into non-placental lineages. In a significant paradigm shift, we now demonstrate that multipotent *CDX2*-expressing cells persist in the cytotrophoblast layer of the human placenta through term. This finding aligns with earlier studies on early-gestation placenta^51^ and highlights the cytotrophoblasts as proliferative, stem-like cells with known migratory and invasive roles during placentation^52^. Notably, *CDX2^+^* cells isolated from term human placentas expressed villous trophoblast markers, displayed clonal proliferation, and showed directed migration—demonstrating key stem/progenitor traits relevant to cardiac regenerative therapies.

To investigate the functional potential of *CDX2*-expressing cells, we employed a lentivirus-based approach to isolate *CDX2^+^* (CDX2mCherry) and mCherry negative cell populations from human placentas. Despite sourcing all samples from healthy term placentas, we observed some variability in proliferative capacity across donors. To account for biological diversity and enhance the generalizability of our findings, we analyzed a large and racially diverse cohort comprising 180 placentas. This comprehensive sampling strategy allowed for a more inclusive assessment of placental progenitor cell function across populations. Isolating CDX2mCherry cells also enabled direct comparison with mCherry negative cells from the same chorionic region. Although the mCherry negative population retained some clonogenic capacity, CDX2mCherry cells exhibited significantly greater cardiovascular differentiation potential. Transcriptomic profiling revealed a marked reduction in mesodermal lineage commitment in mCherry negative cells, which corresponded to minimal cardiomyocyte development (**Extended Data Fig. S5, S6a–d**, **Fig. 5**). This likely reflects, in part, the incomplete transduction of all *CDX2*-positive cells by the lentiviral system—resulting in a small subset of true *CDX2^+^* cells within the mCherry negative population, a known limitation of our approach. Nonetheless, compared with both mCherry negative cells and hESC controls, CDX2mCherry cells showed significantly increased expression of genes associated with chemotaxis, migration, and cardiovascular differentiation.

Consistent with transcriptomic analyses, human CDX2mCherry cells rapidly gave rise to spontaneously beating cardiomyocyte clusters within five days of co-culture on neonatal murine cardiomyocyte feeders. By day 10, approximately 70% of CDX2mCherry cells had differentiated into cardiomyocytes. To exclude the possibility of cell fusion with feeder cardiomyocytes, we performed XY fluorescence in situ hybridization (FISH), confirming diploid human nuclei within CDX2mCherry cells. Notably, the onset of beating occurred significantly earlier than in conventional co-culture models or directed differentiation protocols using pluripotent stem cells, which typically require 8–12 days to initiate contractility^31,53-55^. This accelerated cardiogenic response likely reflects the enriched pre-primed cardiogenic transcriptome intrinsic to CDX2mCherry cells. These findings suggest that CDX2mCherry cells represent a promising progenitor population for efficient, rapid cardiomyocyte generation with translational relevance for cardiac regenerative therapies.

To validate these observations in a defined, feeder-free system, we subjected CDX2mCherry cells to a Wnt-modulated cardiac differentiation protocol. This approach yielded robust populations of cTnT^+^ cardiomyocytes (**Fig. 4**), with significantly higher frequencies of differentiated cells observed as early as day 8, compared to H9ESCs. By day 20, CDX2mCherry cells exhibited a markedly higher overall yield of cardiomyocytes. While we observed sample-level heterogeneity-including variable retention or downregulation of mCherry expression across placental *CDX2* cell populations, CDX2mCherry cells consistently outperformed hESCs in both the kinetics and efficiency of cardiomyocyte differentiation under standardized conditions^31,54^. These results underscore the unique regenerative potential of CDX2mCherry cells and position them as a compelling candidate for cell-based cardiac repair.

To assess the therapeutic potential of CDX2mCherry cells in cardiac repair, we administered them to NOD/SCID mice subjected to experimental MI. Functional assessment by cardiac MRI revealed a significant improvement in left ventricular function in mice receiving CDX2mCherry cells, compared to both mCherry negative cell recipients and vehicle-only controls, which showed progressive functional decline one month post-MI. Histological analysis further demonstrated enhanced *in vivo* cardiomyocyte differentiation and markedly reduced myocardial fibrosis in the CDX2mCherry group. Due to the xenogeneic nature of the human cells, we utilized immunodeficient NOD/SCID mice. However, we observed that this strain did not tolerate large infarcts as well as previously used models^32^ necessitating a modified MI protocol to induce small-to-moderate infarcts with baseline post-MI ejection fractions of ∼35–40%.

For proof-of-concept delivery, CDX2mCherry cells were injected intramyocardially. This approach was selected because the relatively large size of the human placental cells precluded efficient intravenous delivery through the mouse tail vein, unlike in prior murine-to-murine transfer studies^30^. Future studies in large animal models will be essential to evaluate intravenous or intracoronary routes of administration, which may be more clinically relevant and better suited to accommodate human cell size. A further limitation of the current model is that the use of immunodeficient mice precludes investigation of the immunomodulatory properties of CDX2mCherry cells in the context of myocardial injury - an important axis of their therapeutic potential that warrants exploration in future studies.

Given the placenta’s progressive vascularization throughout gestation, placental cells are inherently primed for angiogenesis. Consistent with this, transcriptomic profiling and *in vitro* assays revealed robust angiogenic potential in CDX2mCherry cells. These cells readily formed endothelial tubes and demonstrated functional uptake of acetylated low-density lipoprotein (Ac-LDL), confirming differentiation into functional endothelial cells. To evaluate their capacity for neo-angiogenesis in a non-injury context, we performed an *in vivo* Matrigel plug assay. CDX2mCherry cells significantly increased vessel density within the plugs, although vessel area was not significantly different compared to plugs seeded with mCherry negative cells. These results underscore the angiogenic competence of CDX2mCherry cells—an essential attribute for therapeutic efficacy in cardiac repair, where successful engraftment and survival of transplanted cells depend heavily on the formation of new vasculature. Interestingly, angiogenesis and the invasive nature of trophoblast cells are interlinked as the migration potential of trophoblast cells directs endothelialization/angiogenesis during placentation ^56-58^. On the other hand, *SDF1*-mediated chemotaxis is further involved in stem cell-mediated homing for regeneration ^58,59^ and CDX2mCherry cells demonstrated significantly higher *SDF1*-mediated migration compared to our two control populations.

Once we established the superior cardiovascular differentiation potential of CDX2mCherry cells—surpassing both hESCs and mCherry negative placental cells by bulk RNA sequencing— we next interrogated the proliferation and lineage specification kinetics of these cells at single-cell resolution. This analysis aimed to determine whether key cardiogenic transcriptional programs were preserved following *in vitro* expansion, and to begin addressing the inherent heterogeneity of placental cell populations. Single-cell RNA sequencing of freshly isolated and expanded CDX2mCherry cells revealed 19 transcriptionally distinct subpopulations, with no evidence of *CDX2* downregulation post-expansion, affirming phenotypic stability.

Among these clusters, we identified subsets with discrete lineage biases: STB populations enriched for cardiogenic gene signatures, and EVT clusters with more vasculogenic profiles. Notably, we observed no significant transcriptomic divergence in angiogenic or cardiogenic gene programs between fresh and expanded CDX2mCherry cells, underscoring their preserved therapeutic potential. Within the cardiogenic compartment, STB2 emerged as a dominant cluster, highly enriched in core cardiac mesoderm regulators—including *NKX2-5*, *IRX3*, *MSX1*, *GATA4*, *BMP4*, *HAND2*, *DKK1*, *SFRP1*, and *FGF2*. *FGF2*, in particular, plays dual roles in promoting angiogenesis and vascular stability via crosstalk with maternal tissues, while also supporting the proliferation and survival of cardiac progenitors during development through interactions with Wnt and BMP pathways^60,61^.

In addition to lineage-specific genes, our scRNA-seq data also revealed the expression of immunomodulatory and chemotactic genes across multiple clusters, with no significant variation post-expansion—suggesting potential roles in immune regulation and homing to injury sites. Interestingly, we identified new surface markers such as *AQP1*, a membrane channel protein implicated in ion transport and cell migration^62^ which was selectively elevated in specific STB subsets. Comparative transcriptomic mapping of CDX2mCherry cells to existing scRNA-seq datasets from iPSCs and hESCs further confirmed that CDX2mCherry cells represent a distinct progenitor population. While these cells align along established differentiation trajectories of pluripotent stem cells toward endothelial and cardiomyocyte fates, they possess a transcriptional identity that is neither redundant with, nor fully recapitulated by, traditional pluripotent systems.

Taken together, these findings provide compelling evidence of the functional competence and developmental versatility of placental CDX2 cells. From their ability to generate spontaneously beating cardiomyocytes and functional endothelial cells *in vitro*, to their *in vivo* capacity to enhance cardiac function and reduce fibrosis post-infarction, placental CDX2 cells represent a new, highly promising progenitor cell population. Importantly, they are derived from a readily available and ethically non-contentious source - term human placenta - offering an abundant reservoir for clinical translation. The diverse regenerative properties of placental CDX2 cells lay the foundation for next-generation, precision-guided cell therapies for cardiac repair and broader vascular regeneration strategies.

## Supporting information

Supplemental tables and guide

## Supplementary Information

Extended Data Figure 1 to 10

Supplementary Tables 1 to 3

Supplemental Movies S1 to S12

## Source data

Source Data Fig. 1 - Sanger sequencing, PEAKON and primer sequence data.

Source Data Fig. 2a - Excel data DEG bulk RNAseq CDX2 versus H9 ESCs.

Source Data Fig. 2b - Excel data DEG bulk RNAseq CDX2 versus mCherry negative cells.

Source Data Fig. 6a - Excel data scRNA-seq-cluster based DEG of CDX2mCherry cells.

Source Data Fig. 6b - Excel data scRNA-seq cardiogenic, angiovascular and immunomodulatory genes.

Source Data Fig. 6c - Excel data scRNA-seq of CDX2mCherry cells, iPSC and human ES comparison.

