## Supplementary material for "A multipotent cell type from term human placenta": Data S1

### Sanger Sequencing of human CDX2 PCR product- FASTA sequence

CTGTACCCTAGCTCCGTGCGCCACTCTGGCGGCCTCAACCTGGCGCCGCAGAACTTCGTCAGCCCCC
CGCAGTACCCGGACTACGGCGGTTACCACGTGGCGGCCGCAGCTGCAGCGGCAGCGAACTTGGACAGCGCGC
AGTCCCCG
GGGCCATCCTGGCCGGCAGCGTATGGCGCCCCACTCCGGGAGGACTGGAATGGCTACGCGCCCGGAGGCGC
CGCGGCCGC
CGCCAACGCCGTGGCTCACGGCCTCAACGGTGGCTCCCCGGCCGCAGCCATGGGCTACAGCAGCCCCGCAGA
CTACCATC
CGCACCACCACCCGCATCACCACCCGCACCACCCGGCCGCCGCGCCTTCCTGCGCTTCTGGGCTGCTGCAAAC
GCTCAAC
CCCGGCCCTCCTGGGCCCGCCGCCACCGCTGCCGCCGAGCAGCTGTCTCCCGGCGGCCAGCGGCGGAACCT
GTGCGAGTG
GATGCGGAAGCCGGCGCAGCAGTCCCTCGGCAGCCAAGTGAAAACCAGGACGAAAGACAAATATCGAGTGGTG
TACACGG
ACCACCAGCGGCTGGAGCTGGAGAAGGAGTTTCACTACAGTCGCTACATCACCATCCGGAGGAAAGCCGAGCT
AGCCGCC
ACGCTGGGGCTCTCTGAGAGGCAGGTTAAAATCTGGTTTCAGAACCGCAGAGCAAAGGAGAGGAAAATCAACAA
GAAGAA
GTTGCAGCAGCAACAGCAGCAGCAGCCACCACAGCCGCCTCCGCCCCTCCCCAGCNTTCAGCC

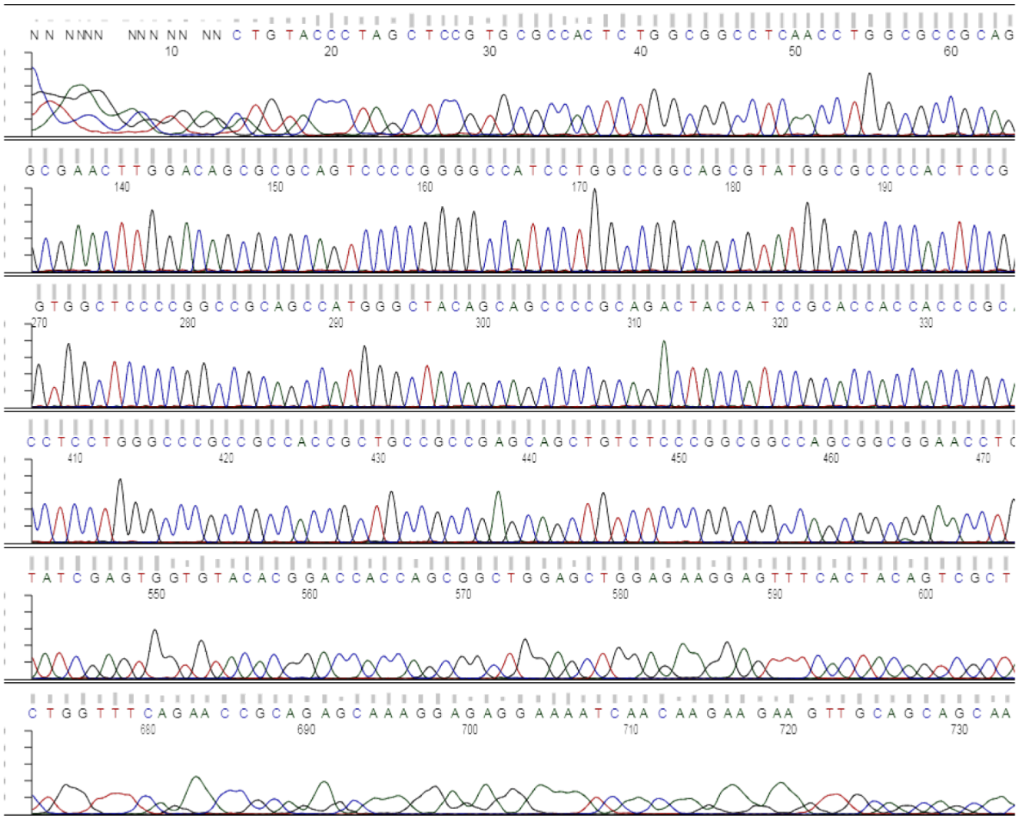

| | |
|---|---|
| Sequence File: | Human_CDX2-hu_CDX2_Forward_primer.seq |
| Average Signal Intensity | G: 690, A: 411, T: 315, C: 662 |

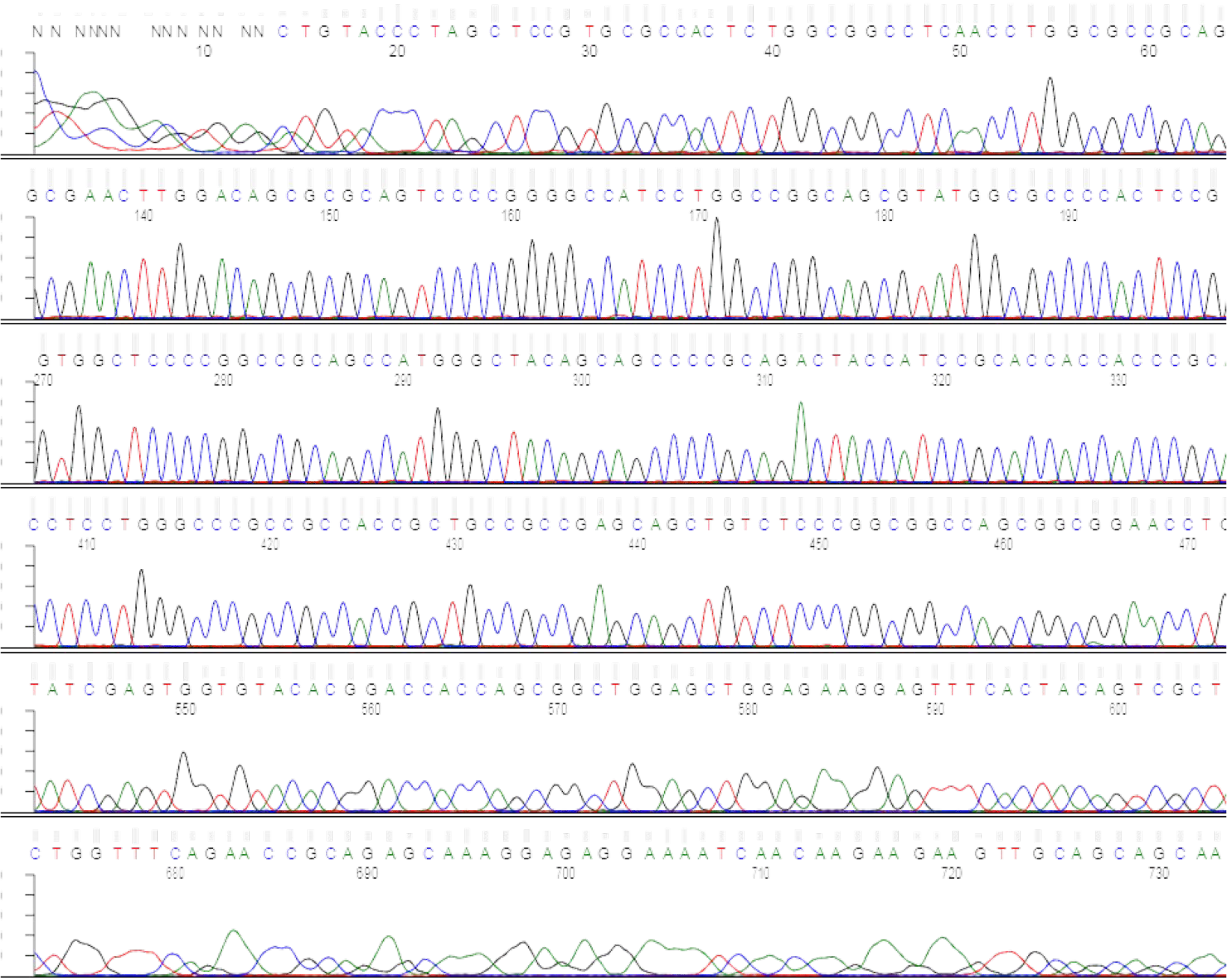


| Sequence File: | **Human_CDX2-hu_CDX2_Forward_primer.seq** |
|---|---|
| Average Signal Intensity | G: **690**, A: **411**, T: **315**, C: **662** |

### Supplemental Materials
### CDX2 promoters

#### A. Human CDX2 promoter #1, 612bps

ATGCATAAACAACCACTGCTCCTGTCTCCAAGCTCAGATTCCTACCAAGATAGCCTTTTCTCTTCCCCT
CTCTTTTGTAAGTCTCTTGATTTCATTCTTTGAACCTGTGATTGGAGGTTAAAGTGCACCAGGTTGGAA
GGAGGAAGCTCTTAACAATAAAGGTTTGAATATTTAGCTGTGTCAGGTCGCTGCCCTCTCACGAGCCTC
CCTCCCCTTTATCTTTTAAAATGCAAATTATGTTTCGAGGGGTTGTGCGTAGAGTGCGCGCTGCGCCTC
GACGTCTCCAACCATTGGTGTCTGTGTCATTACTAATAGAGTCTTGTAAACACTCGTTAATCACGGAAG
GCCGCCGGCCTGGGGCTCCGCACGCCAGCCTGTGGCGGGTCTTCCCCGCCTCTGCAGCCTAGTGGGAAG
GAGGTGGGAGGAAAGAAGGAAGAAAGGGAGGGAGGGAGGAGGCAGGCCAGAGGGAGGGACCGCCTCGGA
GGCAGAAGAGCCGCGAGGAGCCAGCGGAGCACCGCGGGCTGGGGCGCAGCCACCCGCCGCTCCTCGAGT
CCCCTCGCCCCTTTCCCTTCGTGCCCCCCGGCAGCCTCCAGCGTCGGTCCCCAGGCAGC

Sequence obtained from manually upstream of Exon1 at ~300bps and ~308bps into Exon1 right before the first ORF ATG SITE

| Position | Score | Likelihood |
|---|---|---|
| 400 | 0.651 | Marginal prediction |

#### B. Human CDX2 promoter #2, 1,173bps

CCTAGGTAAGCATTAGCAGAAATTCTCTTTTCCTTATATTTAAGTATAATAAACATACAAGTGTAGCTCAATGAATTTTCACA
AACTGACATTCTGTGTAACCAGCAGCCTAAGAAACTGCTTTACCAACGATCCCCTAGCTCGCCTCCAGTTATGCACGCCAATA
ACCACTAGCCTAACTTCTACCACATGCCCATTACTTCTGTAGTTTAAAACTTCTGATTCTTGAATGTAAACGTTTAACAATAA
ATCGCTTGAATTTAACTCAAATTTCAAATGTAAGATGAAGTCAGAGATGCAGCCTGAATCTAGGATCATAATTTGTCTTGTGC
GGAGGGCGAGTAATTTCCTTGGGCAAGAAAATAACTGGAGGTGACAGTTGTTTGGGGCTGCAGTCGTCCGGGCCAGGAGCACA
GGGCGGGAAGGAATGGCCCATCTCTTAGGGCTCTCTGCTTGTCACCTACCAGGTTGGTCAGAAACGTTCTCATCAAAGCAATG
GTTCTCTTTTCTTTTCTCTTTGGGACAGAAGGAGTTTCTTGACCGCCCTCTTCCCTGCAAATGCATAAACAACCACTGCTCCT
GTCTCCAAGCTCAGATTCCTACCAAGATAGCCTTTTCTCTTCCCCTCTCTTTTGTAAGTCTCTTGATTTCATTCTTTGAACCT
GTGATTGGAGGTTAAAGTGCACCAGGTTGGAAGGAGGAAGCTCTTAACAATAAAGGTTTGAATATTTAGCTGTGTCAGGTCGC
TGCCCTCTCACGAGCCTCCCTCCCCTTTATCTTTTAAAATGCAAATTATGTTTCGAGGGGTTGTGCGTAGAGTGCGCGCTGCG
CCTCGACGTCTCCAACCATTGGTGTCTGTGTCATTACTAATAGAGTCTTGTAAACACTCGTTAATCACGGAAGGCCGCCGGCC
TGGGGCTCCGCACGCCAGCCTGTGGCGGGTCTTCCCCGCCTCTGCAGCCTAGTGGGAAGGAGGTGGGAGGAAAGAAGGAAGAA
AGGGAGGGAGGGAGGAGGCAGGCCAGAGGGAGGGACCGCCTCGGAGGCAGAAGAGCCGCGAGGAGCCAGCGGAGCACCGCGGG
CTGGGGCGCAGCCACCCGCCGCTCCTCGAGTCCCCTCGCCCCTTTCCCTTCGTGCCCCCCGGCAGCCTCCAGCGTCGGTCCCC
AGGCAGCATGG

GenBank: EU544225.1

>EU544225.1 Homo sapiens caudal type homeobox 2 (CDX2) gene, promoter region and 5' UTR

| Position | Score | Likelihood |
|---|---|---|
| 900 | 0.651 | Marginal prediction |

#### C. Human CDX2 promoter #3, 1,917bps

GGAACCAGAAAACAGGGGATCCCGCAGCCCTAGGCTAGTTCTGATCGCTTTCAGGTGTCTGCAGAGGCAAGTTGCTGGTTGTC
ACCTGTAAAATGGGGAGGATAAAAACACCTCCCAGATTTTGTTCTAGATCCTAGGGGGATGTGAGGCTCAAGGGAGATAAAGG
ACACTGGAGAGCACCCTAGAAATGACAGGATGAAGGCGATGGTGACAAATATCCGAGCGAAACGCTTGACAATGAGAACAGAC
AAGTGCAGGTCTCCAGGAGTGCCGCGAGCGCCCGCGGGTTCTGAGAGCGCTCAAAGCCGCCGAGTCAGGCTGCCCAGCCCGCC
GGGCCTCGCCGCAGTGATCCTCATTCCCGAATCTGGCAGCGCTGTCAAAGGCTTGTATTAGGAGGTGAACGGCGGCCGCAGGC
CCACTCCACGCGGTTGCTGAAACCGAGCTGGGCGCGCGGGGGCCGAATCTCGCCGCCTCCGCGCTCCTGTCGGGGCAGCTC
CCGATCCCGGGCTGCGCGGCTTCGGTCCCCAAGACGGCCACTTCCAGCCCTAGGCCCCTTGGCCGCAGCGCTTCCCAAACCAA
GAGAGATCCTTTCTCAACTCAGAGCTTTTCATTAGCAGTCGTTAATAATGGCCCTGAGTTGCCTTATCATCTCCTGGAAATGA
GAAATAAATTTCTTCGGAGAACGTTTCCCTTTGTAAAGGACAGAGAGTTTTAAAGATACAGGTATGATGTAAGACACATAAAT
ACCTAGGTAAGCATTAGCAGAAATTCTCTTTTCCTTATATTTAAGTATAATAAACATACAAGTGTAGCTCAATGAATTTTCAC

AAACTGACATTCTGTGTAACCAGCAGCCTAAGAAACTGCTTTACCAACGATCCCCTAGCTCGCCTCCAGTTATGCACGCCAAT
AACCACTAGCCTAACTTCTACCACATGCCCATTACTTCTGTAGTTTAAAACTTCTGATTCTTGAATGTAAACGTTTAACAATA
AATCGCTTGAATTTAACTCAAATTTCAAATGTAAGATGAAGTCAGAGATGCAGCCTGAATCTAGGATCATAATTTGTCTTGTG
CGGAGGGCGAGTAATTTCCTTGGGCAAGAAAATAACTGGAGGTGACAGTTGTTTGGGGCTGCAGTCGTCCGGGCCAGGAGCAC
AGGGCGGGAAGGAATGGCCCATCTCTTAGGGCTCTCTGCTTGTCACCTACCAGGTTGGTCAGAAACGTTCTCATCAAAGCAAT
GGTTCTCTTTTCTTTTCTCTTTGGGACAGAAGGAGTTTCTTGACCGCCCTCTTCCCTGCAAATGCATAAACAACCACTGCTCC
TGTCTCCAAGCTCAGATTCCTACCAAGATAGCCTTTTCTCTTCCCCTCTCTTTTGTAAGTCTCTTGATTTCATTCTTTGAACC
TGTGATTGGAGGTTAAAGTGCACCAGGTTGGAAGGAGGAAGCTCTTAACAATAAAGGTTTGAATATTTAGCTGTGTCAGGTCG
CTGCCCTCTCACGAGCCTCCCTCCCCTTTATCTTTTAAAATGCAAATTATGTTTCGAGGGGTTGTGCGTAGAGTGCGCGCTGC
GCCTCGACGTCTCCAACCATTGGTGTCTGTGTCATTACTAATAGAGTCTTGTAAACACTCGTTAATCACGGAAGGCCGCCGGC
CTGGGGCTCCGCACGCCAGCCTGTGGCGGGTCTTCCCCGCCTCTGCAGCCTAGTGGGAAGGAGGTGGGAGGAAAGAAGGAAGA
AAGGGAGGGAGGGAGGAGGCAGGCCAGAGGGAGGGACCGCCTCGGAGGCAGAAGAGCCGCGAGGAGCCAGCGGAGCACCGCGG
GCTGGGGCGCAGCCACCCGCCGCTCCTCGAGTCCCCTCGCCCCTTTCCCTTCGTGCCCCCCGGCAGCCTCCAGCGTCGGTCCC
CAGGCAGC

| **Position** | **Score** | **Likelihood** |
|---|---|---|
| 300 | 0.679 | Marginal prediction |
| 1100 | 1.073 | Highly likely prediction |
| 1700 | 0.651 | Marginal prediction |
