## Supplementary material for "A multipotent cell type from term human placenta": Methods

**Human subjects**

Pregnant women enrolled in this study ranged in age from 18 to 45 years, were ethnically and racially diverse, and reflected the demographics of those served by our Maternal Fetal Medicine Practice. 189 patients are currently enrolled in this study. Placenta samples were collected under two cohorts. 1) Institute-approved IRB protocol (20-0068, IF2646084), which reflects a prospective cohort of 189 enrolled patients (including future due dates), from which 31 samples have been collected thus far, based on the viable delivery window (weekdays, 9:00 am-5:00 pm). Cohort 2, the de-identified cohort, contained 149 fresh tissue samples under the auspices of the Institutional Biobank Pathology Core facility. De-identified samples (all identifying patient information was replaced by a BRP #ID) at the Biorepository Core Facility were sampled by a neonatal placental pathologist. Both sample collections adhered to the standard protocol to prevent interference with any necessary clinical assessment of the placenta. Only 5 g of tissue from the chorionic plate was collected from each placenta immediately after delivery, as the remainder of the placenta would be stored by Pathology for clinical/diagnostic purposes. Unless otherwise specified, each experiment was performed using at least three different patient samples (n = 3).

**Cell line and Primary cultures**

*Cell lines:* The H9 human embryonic cell line (Stem Cell Engineering Core, WA09 (ESC) https://hpscreg.eu/cell-line/WAe009-A) was used as a control for transcriptomics and cardiac differentiation experiments. H9ES cells, HEK 293T cells, and HeLa cells (ATCC) served as negative controls that lack or express very low endogenous CDX2, respectively, while the human colon adenocarcinoma cell line DLD1 (CCL-221 ATCC) served as a positive control for CDX2. RPMI (Roswell Park Memorial Institute, ThermoFisher, Cat#11875119) medium, containing 10% Fetal Bovine Serum (FBS, ThermoFisher, Cat#A5256801), was used as the culture medium for both HeLa and DLD1 cell lines.

*Primary cultures:* Human Umbilical Vein Endothelial Cells (HUVECs) were maintained in M200 endothelial basal medium and supplemented with 1X Low Serum Growth Supplement (LSGS, ThermoFisher, Cat# S100310) during differentiation. Isolated chorionic cells and CDX2mCherry positive and mCherry negative cells were plated using Iscoves Modified Dulbecco’s Medium (IMDM, ThermoFisher, Cat#12440053) +10 % FBS.

**Placental CDX2 cell isolation**

Placentas were received within 30 to 120 minutes after delivery. The gross appearance was briefly examined, including identification of the cord insertion site and disc edges. Approximately a 5.0 x 2.0 cm rectangular area of each full-thickness disc was incised with a surgical blade halfway between the cord insertion site and the disc edge, avoiding obvious sub-chorionic fibrin deposition. Any remaining amnion was removed using forceps. Finally, the decidual surface was shaved with a blade to minimize the maternal component. The chorionic tissue was then quickly transferred to 1X Hanks Balanced Salt Solution (HBSS) containing 2X penicillin and streptomycin and kept on ice. The fresh tissue was thoroughly rinsed with 1X HBSS to remove blood, minced into approximately 1mm³ pieces, and transferred to 50 mL Falcon conical tubes containing 2 mg/mL Collagenase Type 2 (Worthington, NJ, Cat#LS0004194) and 1 mg/mL Pronase (Calbiochem, Cat#53702) in 1X HBSS. The samples were incubated in the enzyme solution for 2 hours at 37°C with 5% CO₂. The tissue mixture containing the enzyme was gently mixed every 20 minutes using disposable pipettes to facilitate digestion. After digestion, the enzyme reaction was halted by adding serum, and the cell mixture was filtered through a 100-micron cell strainer (Falcon, Cat#352360) and centrifuged at 1600 rpm for 4 minutes. Red blood cells were removed using 1X RBC lysis buffer (10X stock, eBioscience, Cat# 00-4300-54) by incubating the cell pellet in the buffer for 15 minutes at room temperature in the dark, followed by centrifugation. The final cell pellet was resuspended in IMDM supplemented with 10% fetal bovine serum (FBS) or the appropriate assay medium for downstream experiments.

**Lentiviral transduction and isolation of placental CDX2 cells**

To select viable CDX2 cells from the total chorionic cells, we designed a lentiviral construct where the human CDX2 promoter drove mCherry fluorescence. The CDX2 gene is oriented in the reverse (3’-5’) direction, and its promoter activity is not as strong as that of traditional upstream promoters. Therefore, we created three different constructs using sequences that span various regions upstream of the 3’ UTR, and in some cases, include a fragment into the start codon to cover promoter region 1 (612 bp), promoter region 2 (1174 bp), and promoter region 3 (1197 bp). Detection was based on mCherry fluorescence, with a ubiquitous CMV promoter in the same vector backbone serving as a control for transduction efficiency. The human adenocarcinoma cell line DLD1 served as a positive control for CDX2 expression. H9ES cells and HEK 293T cells served as negative controls. Based on the optimization experiments **(Extended Data Fig. 3d**), promoter 1 showed the strongest mCherry fluorescence; therefore, we used lentiCDX2 (612 bp) for all subsequent experiments. For transduction, chorionic cells were plated at a density of 2 × 10^5 cells per well in IMDM with 10% FBS and allowed to adhere overnight. The next day, cells were transduced with lentivirus at a multiplicity of infection (MOI) of 50, using 8 µg/mL polybrene to enhance transduction. The plates were spinoculated at 2000 rpm for 1 minute and incubated at 37 °C with 5% CO_2_ for 48–72 hours. After 48 hours, the medium was replaced, and cells were examined for mCherry fluorescence under a Zeiss AxioObserver epifluorescence microscope. CDX2mCherry and mCherry negative populations were sorted via flow cytometry using a BD Influx/IMI5L/Cytek Aurora sorter. With this method (at passage 0), we initially obtained between 2% and 10% CDX2mCherry cells. We then allowed the chorionic cells to reach confluence (replenishing fresh media to remove floating and dead cells on the third day), re-plated, and transduced them (passages 1-3) as described in **Fig. 1e**. This strategy increased transduction efficiency, enabling us to isolate approximately 30% CDX2 mCherry cells from the plated samples **(Fig. 1f).**

**Immunofluorescence**

At the time of collection, small sections from the same chorionic area were fixed in 10% formalin and embedded in paraffin. Alternatively, to detect CDX2 expression, amniotic tissue/membrane was also collected in some samples. Tissue sections (7 µm) were deparaffinized and rehydrated using a sequential series of steps involving xylene, 100% ethanol, 90% ethanol, 80% ethanol, and 70% ethanol, followed by deionized water. Antigen retrieval was achieved by proteolytic-induced epitope retrieval (PIER) using a combination of trypsin and proteinase K for 10 min at room temperature, followed by washing with 0.2% Tween-20. Peroxidase quenching in 3% H2O2 in methanol (10 min) was performed to prevent endogenous peroxidase from cleaving the substrate, followed by washing with ice-cold 1X PBS. The sections were then permeabilized using 0.25% Triton X-100 in 1X HBSS for 20 minutes at room temperature (RT), followed by blocking with 5% Bovine Serum Albumin (BSA) in 1X HBSS for 2 hours. Samples were incubated overnight at 4°C with primary antibodies including: anti-huCDX2 Abcam, Cat# ab86949, anti-cytokeratin-7, Invitrogen Cat# MA5-53274, anti-HLA-G, ThermoFisher, Cat# PA5-98143), anti-vWF ThermoFisher, Cat#98143, and anti-CD31 at a 1:100 dilution. The next day, secondary antibodies (ThermoFisher anti-ms Alexa 568 cat#A11004 anti-ms Alexa 488 cat#A21202,anti-rb Alexa 647, cat# A21244, anti-rbAlexa 488 cat#31635, and anti-rb Alexa 555, Cat#31572 (at 1:200 dilution) were added for 1h at room temperature. Auto fluorescence was then quenched using 0.7% Sudan Black B (#S2380, Sigma-Aldrich) in 70% ethanol for 5 min at room temperature. Slides were then rinsed three times using 1X HBSS, and nuclei were counterstained with DAPI, and cover-slipped with mounting media (KPL, Gaithersburg, MD, USA, or Vectorlabs H-1000). Similarly, after the co-culture experiments, the cells were fixed, permeabilized, blocked, and primed with primary and secondary antibodies. Cells were later fixed, permeabilized, and immunostained using cardiac troponin T (Santa Cruz Biotechnology, Cat#sc-20025) and cardiac sarcomeric actin (Abcam, Cat#ab68168), as described above. For *in vivo* angiogenesis assay and myocardial infarction experiments, the Matrigel and heart sections were removed (for heart sections, after isoflurane anesthesia, hearts were injected with 3M KCL for diastolic arrest, followed by euthanasia and excision of hearts), stored in 10% formalin overnight, followed by paraffin embedding and grossing. 5µ sections were cut from Matrigel blocks and MI hearts onto slides and were deparaffinized using sequential steps as described above and immunostained. For the angiogenesis experiment, rabbit anti-human CD31 was used with endogenous mCherry fluorescence. For post-MI heart sections, anti- cTnT (Cat#ab45932, Cat#sc-20025 Santacruz Inc, MA5-12960 ThermoFisher), mouse anti-human sarcomeric actinin (Cat#ab68168 Abcam), rat CD31 (Cat#sc-101454, SantaCruz Inc, MA1-19199, ThermoFisher), mouse antihuman smooth muscle actin (Cat#ab5694, Abcam), rat anti-mCherry (Cat#M11217, ThermoFisher) Chicken anti-mCherry (Cat#NBP2-25158, Novus Biologicals), goat anti-human Ku80 (Cat#AF5619, R&D), mouse anti-human Ku80 (Cat#611360, BD Biosciences) at 1:100 dilutions were used. Appropriate secondary conjugated antibody dilutions (1:200) were made in Antibody diluent (Dako, Agilent, S080983-2). Secondary antibodies, besides those mentioned above, include Anti-rat Alexa 568 (Cat# A11077, ThermoFisher), anti-mouse Alexa 647 (Cat#A21235, ThermoFisher), and Anti-Chicken 647 (Cat#A78952, ThermoFisher). Sections were mounted as described, and imaged using a Zeiss Observer Z1 inverted fluorescent microscope with Zen Blue software and Zeiss Confocal microscopy (Carl Zeiss, Munich, Germany).

**RNA extraction, cDNA preparation, PCR, and Sanger sequencing**

Chorionic villus samples (CVS) was a kind gift from a colleague at the institution and for transcript analysis, random areas of the fetal chorionic region from three different donors were commercially obtained from Advanced Tissue Services (Phoenix, AZ, USA). Total RNA was extracted using the Direct-zol RNA Miniprep Kit (Zymo Research). The eluted RNA was purified to remove traces of genomic DNA with DNase (DNA-free kit – DNase treatment, Ambion), and the RNA concentration was measured using a Qubit HS RNA assay kit (Thermo Fisher Scientific). A total of 2 μg of RNA was reverse-transcribed into cDNA using the Maxima First Strand cDNA Kit (ThermoFisher), following the manufacturer’s instructions. PCR was performed with human-specific CDX2 primers (Table S1). GAPDH served as the internal control. The PCR cycling conditions included 50°C for 2 minutes (1 cycle), 95°C for 10 minutes (1 cycle), then 40 cycles of 95°C for 15 seconds and 60°C for 1 minute. The results were normalized to the endogenous control genes and analyzed using the comparative Ct method. Specific primers for human CDX2 (Table S1), optimized for sequencing (product size 1.5 kb), were synthesized, and 10-15 μL of crude PCR product was mixed with forward sequencing primers for Sanger sequencing (Genewiz, Azenta, USA). The FASTA output sequences were validated through NCBI nucleotide BLAST (BLASTn) to confirm proper alignment with the human CDX2 transcript.

**Protein extraction and western blot analyses**

Frozen placental tissues, freshly isolated and cryopreserved chorionic samples (pooled n=3), DLD1 cells, HeLa cells, and undifferentiated H9ES cells were thawed on ice and incubated in 1X RIPA buffer (with 1 % SDS and 1X protease inhibitor cocktail, Sigma Aldrich, Cat#539195) for protein extraction. Samples were minced on ice, vortexed every few minutes for a maximum of 45 min, and homogenized using a cordless handheld microtube homogenizer (UX-44444-70, Cole-Parmer, USA), followed by centrifugation at 12,000 rpm at 4°C for 15 min, and the supernatant (protein lysate) was transferred to another microcentrifuge tube. Protein samples were stored at -80 °C and/or quantified using a Qubit® 3.0 Fluorometer (ThermoFisher Scientific, MA, USA) using the Qubit Protein Assay Kit and also BCA-based protein quantification in the nanodrop. Identical concentrations of protein lysates (approximately 80µg, **see Fig. 1b**) were loaded along with 5X Laemmli buffer in a 10% Mini-Protean TGX Precast Protein Gel, 10-well, 30 µL (Bio-Rad, Hercules, CA). The gel was run at 80 V and then at 100 V in 1X Tris-Glycine-SDS Running Buffer (Tris base, 250 mM; glycine, 1.92 M Boston BioProducts, Ashland, MA, Cat#BB-2689). Pre-stained SDS-PAGE standard protein markers (Bio-Rad, Hercules, CA [Cat# 161-0318]) and biotinylated protein ladder (Cell Signaling Technology, Inc., Danvers, MA, Cat# 7727) were used as controls. Following gel electrophoresis, proteins were transferred to a PVDF membrane in 1X Transfer Buffer (Tris base, 0.25 M, glycine, 1.92 M) (Boston Bioproducts, Ashland, MA) overnight at 14 V. The PVDF membrane was blocked with 5% BSA (MP Biomedicals, LLC, Solon, OH) in 1X Tris-Buffered Saline (Boston BioProducts, Ashland, MA) supplemented with 0.05% Tween 20 (TBST) (Thermo Fisher Scientific, MA, USA), followed by 2.5 % dry milk in TBST for 2h. The rabbit CDX2 primary antibody #Cat 3977 (Cell Signaling Technology, Inc., Danvers, MA, USA) was used at a dilution of 1:800. Cell Signaling HRP-linked anti-rabbit IgG secondary antibody #Cat 7074 was used at a dilution of 1:1000. Anti-biotin HRP-linked secondary antibody #Cat 7075 was used at a dilution of 1:3000 (Cell Signaling Technology, Inc, USA). GAPDH #Cat 97166 (loading control) antibody was used at a dilution of 1:1000 with anti-mouse HRP antibody #Cat 7076 (Cell Signaling Technology, Inc., USA).

**Ultralow-input Bulk RNA sequencing**

Flow cytometry-based sorting was used to isolate CDX2-expressing cells, identified by their mCherry fluorescence, from the chorionic cell population of de-identified placental samples (n = 5). Isolated CDX2mCherry cells were centrifuged, and 200 μL Tri reagent (ThermoFisher Scientific, USA) was added and incubated at room temperature for 15 minutes. The samples were stored at -80°C until the RNA extraction and library preparation were performed. The library preparation workflow is depicted below for sequencing on Illumina Hiseq (HiSeq™ 2000 Sequencing System, Illumina, Genewiz/Azenta Life Sciences).

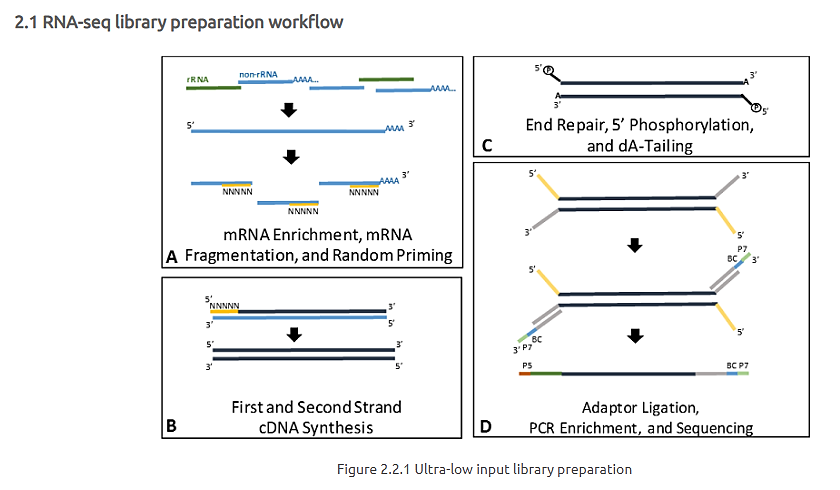

Ultra-low input RNA seq library preparation workflow

*Distribution of read counts*

The distribution of read counts in libraries were examined before and after normalization. The original read counts were normalized to adjust for various factors such as variations of sequencing yield between samples. These normalized read counts were used to accurately determine differentially expressed genes.

*Mapping the sequence read to the reference genome*

Sequence reads were trimmed to remove possible adapter sequences and nucleotides with poor quality using Trimmomatic v.0.36. The trimmed reads were mapped to the Homo sapiens GRCh38 reference genome available on ENSEMBL using the STAR aligner v.2.5.2b. The STAR aligner is a splice aligner that detects splice junctions and incorporates them to help align the entire read sequences. BAM files were generated as a result of this step.

*Sample similarity assessments*

Data quality assessments were performed to detect any samples that are not representative of their group, and thus, may affect the quality of the analysis. The overall similarity among samples were assessed by the euclidean distance between samples. This method was used to examine which samples are similar/different to each other and if they fit to the expectation from the experiment design

*Extracting gene hit counts*

Unique gene hit counts were calculated by using featureCounts from the Subread package v.1.5.2. The hit counts were summarized and reported using the gene_id feature in the annotation file. Only unique reads that fell within exon regions were counted. If a strand-specific library preparation was performed, the reads were counted strand-specifically.

*Differential gene expression analyses*

After the extraction of gene hit counts, the gene hit counts table was used for downstream differential expression analysis. Using DESeq2, a comparison of gene expression between the CDX2mCherry positive cells and h9ES cell control (Group 1) and CDX2mCherry cells and mCherry negative cells (Group 2) was performed. The Wald test was used to generate p-values and log2 fold changes. Genes with an adjusted p-value < 0.05 and absolute log2 fold change > 1 were called differentially expressed genes for each comparison.

*Gene ontology analyses*

A Gene Ontology analysis was performed on the statistically significant set of genes using GeneSCF v.1.1-p2. The goa_human GO list was used to cluster the set of genes based on their biological processes and determine their statistical significance. A list of genes was generated, clustered based on their gene ontologies.

**Single-cell RNA sequencing**

*Sample QC and Library preparation*

Human CDX2mCherry cell samples, both freshly isolated (n=2) and expanded at passage 3 (n=3), were thawed for analysis (total n=5). Cell counts were assessed using the Cellaca Cell Counter (Nexcelom Bioscience, Lawrence, MA, USA) in conjunction with NucBlue for staining cell nuclei and propidium iodide (PI) for identifying dead cells. Samples were then processed with the Chromium X platform (10x Genomics, Pleasanton, CA, USA), following the manufacturer’s protocols, to target the capture of approximately 3,000 cells per sample for downstream analysis. Single-cell RNA-seq libraries were prepared using the Chromium Next GEM Single Cell 3’ Gene Expression Kit. Library quality was assessed using the Agilent TapeStation (Agilent Technologies, Palo Alto, CA, USA), and quantified via Qubit 2.0 Fluorometer (Invitrogen, Carlsbad, CA, USA). Additional quantification was performed using quantitative PCR (qPCR) (Applied Biosystems, Carlsbad, CA, USA) prior to sequencing on an Illumina platform, following 10x Genomics' recommended sequencing configurations.

*Data Processing and Analysis*

Raw sequencing data, in the form of .bcl files generated by Illumina, were converted into FASTQ files and de-multiplexed using the 10x Genomics Cell Ranger mkfastq pipeline. QC was carried out by removing doublets from analysis, excluding cells that showed >8,000 feature counts (potential multiplets) or mitochondrial genes at > 50% or hemoglobin genes at >5%. Differential gene expression analysis was conducted interactively using the 10x Genomics Loupe Browser for data visualization and interpretation. Cell annotation was performed using cell marker identification based on classical trophoblast gene markers and the upregulated gene expression profiles from our dataset. Cell markers are displayed as plots for each cell population in either the figure legend or the supplementary material.

**Pseudotime trajectory analyses**

*Data Processing and Quality Control*

A total of 19,594 filtered CDX2mCherry cells were analyzed in this study. Additionally, we used previously published scRNA-seq datasets^33-35^ as follows: iPSC day_0, iPSC-CM day_5, and iPSC-CM day_30 (accession no: E-MTAB-6268) iPS-EC day_8 and iPS-EC day_12 (*2*) (GSE116555). Each replicate/sample within these conditions underwent independent processing, with the following quality control filters consistently applied across all datasets: (i) Cells with more than 8,000 detected genes or more than 50% mitochondrial gene expression were excluded from analysis. (ii) Only genes expressed in at least 10 cells across the dataset were retained for downstream analysis. We also used the undifferentiated ES cell dataset GSM5537050 (3) and the iPSC dataset, and compared them to the CDX2mCherry cells in a comparison heatmap. Differentially expressed genes (DEGs) were identified in the CDX2mCherry cells, iPSC, and ESC populations for cell-type marker genes. The MAST method in Seurat was used for DEG analysis.

*Data integration, dimensionality reduction, and clustering*

Following quality control, datasets from the CDX2mCherry cells and other datasets were merged. RPCA-based integration was performed using the Seurat package (v5.1.0) with the following parameters: normalization. method = "SCT"; dims = 1:30. This integration method aligns gene expression profiles across different conditions while minimizing batch effects. Following integration, dimensionality reduction was performed using UMAP-based clustering. A resolution parameter of 0.5 was applied to identify distinct cell populations within the integrated data. To infer the developmental trajectories of cells, pseudo-time analysis was performed using the Monocle3 package (v1.3.7)*.* Cells were ordered along a trajectory inferred from the UMAP-based clustering, with the CDX2mCherry cells near day 0 iPSCs designated as the starting point. The trajectory was constructed using the learn_graph function, and pseudo-time was computed for each cell along this trajectory. Results were visualized on UMAP plots, with cells colored according to their pseudo-time values.

**Clonality assay**

FACS isolated CDX2mCherry cells were plated at a density of 1-5 cells per well in 60-well Terasaki plates (Thermo Scientific, Cat#163118) in 15µl of growth medium (IMDM supplemented with 10% FBS) and incubated in a humidified carrier plate (150mm cell culture dish). Fresh growth medium was replenished once at day 7. One day after plating, each well was individually imaged to exclude wells with fewer than one or more than one cell. Colonies were counted and imaged on day 14. A colony is defined as a cluster of more than 100 cells, with approximately more than 50% confluency per well.

**Transwell migration assay**

Cell-permeable, 24-well-compatible 8 µm PET inserts (ThermoFisher) were used for migration experiments. IMDM + 2% FBS was added to the lower well of 24-well plates, followed by 100 ng of human SDF-α (R&D), excluding some wells to detect any spontaneous migration. Inserts were carefully placed in each well. In some wells, 1µM AMD3100 (a CXCR4-blocking peptide) was added. An equal number of CDX2mCherry and mCherry negative cells were added to the upper Transwell chamber and allowed to migrate in response to the chemotactic signals from SDF-1 below. There were two biological replicates for each experiment, and three different placenta samples from three patients were analyzed. The plates were incubated at 37°C for 3 hours. After that, each transwell was carefully removed, and the cells remaining in the transwell (non-migrated) and the bottom well (migrated) were collected and manually counted. Migration was quantified using the equation: percent migration = (migrated cells / Total cells) × 100. The data was analyzed and graphically plotted using GraphPad Prism software version 10.

**Endothelial tube formation assay**

Functional angiogenesis was assessed using a tube formation assay. Growth factor-reduced Matrigel was used as the matrix (Corning Inc., USA, Cat# 356230). Matrigel was thawed at 4°C overnight, and while placed on a cooling rack, 100 μL of Matrigel was carefully dispensed into a 96-well plate. The plates were incubated at 37°C for one hour to promote gel polymerization. CDX2mCherry cells were cultured in IMDM+10% FBS until reaching confluence (3-4 days), then trypsinized, and 2 × 10^4 cells were plated per well on Matrigel with M200 medium (GIBCO-human large vessel endothelial cell basal medium) supplemented with Low Serum Growth Supplement (LSGS). LSGS (Thermo Fisher, USA) contained 2% v/v fetal bovine serum, 1 μg/mL hydrocortisone, 10 ng/mL human epidermal growth factor, 3 ng/mL basic fibroblast growth factor, and 10 μg/mL heparin. Human Umbilical Vein Endothelial Cells (HUVEC) (positive control), unfractionated chorionic cells, CDX2mCherry cells, and mCherry negative cells were plated using the same supplemented M200 medium. The plates were incubated at 37°C with 5% CO2 for six hours; images were taken at both 3- and 6-hour time points. Tube formation was quantified using the ImageJ analysis tool, incorporating the Angiogenesis Analyzer plugin.

**Acetylated Low-Density Lipoprotein (AcLDL) uptake assay**

Uptake of AcLDL by CDX2-derived endothelial cells (ECs) was also evaluated. Cells were cultured with 10 ng/mL Vascular Endothelial Growth Factor (VEGF, Peprotech, Cat#100-20) for 7 days in the M200 medium. The medium was then removed, and cells were incubated with 10 μg/mL Alexa Fluor 488-conjugated AcLDL (ThermoFisher, Cat#L23380) in 1X HBSS at 37°C with 5% CO2 for 4 hours. After incubation, the medium was aspirated, and cells were fixed in 4% (v/v) paraformaldehyde in PBS for 30 minutes at 4°C in the dark. LDL uptake was analyzed by examining Alexa Fluor 488 fluorescence using a Leica DM IL LED Inverted Microscope with LAS X imaging multichannel acquisition software (Leica Biosystems, USA).

**Matrigel plug assay for *in vivo* angiogenesis evaluation**

The Matrigel plug angiogenesis assay was adapted from a previously described method (Kastana, P et al., Methods Mol Bio 2019). Briefly, NOD/SCID mice were injected subcutaneously in both flanks with 500 µl of Matrigel supplemented with VEGF (10 µg/ml stock and 100 ng/500 µl working concentration), with or without the addition of HUVEC cells, CDX2mCherry, and mCherry negative cells. (Matrigel (n=2); Matrigel + VEGF (n=2); Matrigel +VEGF+HUVEC (n=2); Matrigel+VEGF+CDX2mCherry cells (n=3); Matrigel+VEGF+mCherry-negative cells (n=3). Fourteen days post-injection, the Matrigel plugs were excised, and the angiogenic responses were initially assessed visually. Subsequent analyses included biochemical and microscopic evaluations. Immunostaining for CD31 was performed to detect endothelial cells. Hematoxylin and Eosin (H&E)-stained, paraffin-embedded sections were imaged using a NanoZoomer S210 Digital Slide Scanner at ×20 magnification. The images were analyzed with NanoZoomer Digital Pathology Image (NDPI) software to assess angiogenesis and neovascularization. Vessel density was quantified by counting the number of vessels in different regions of the plug, and the area occupied by vessels was measured relative to the total plug area.

**Cardiomyogenic differentiation of CDX2mCherry cells in neonatal cardiomyocyte feeder**

Glass-bottom 96-well plates were coated with 1 µg/ml laminin (Corning, Cat#354232, USA). Neonatal cardiomyocyte feeder layers were prepared and plated for 4-5 days until they began beating. Freshly sorted CDX2 mCherry cells were then plated at a density of 1-10^4 to 2x10^4 cells/well onto these contracting myocyte feeders and cultured using IMDM and 10% FBS. Cells were monitored for over four weeks, and live cell imaging was performed to capture spontaneous beating using a Zeiss Axiovision Observer Z1 inverted fluorescent microscope equipped with Axiovision software (Carl Zeiss, Munich, Germany). Cells were later fixed, permeabilized, and immunostained for cardiac troponin T (Santa Cruz Biotechnology, Cat#sc-20025), cardiac sarcomeric actin (Abcam, Cat#ab68168), and endothelial markers (CD31, ThermoFisher, Cat#14-0319-82; vWF, BD Pharmingen, Cat#555849) to detect cardiomyocytes and endothelial cells derived from human CDX2mCherry cells.

**XY chromosome analysis**

Mouse XY chromosome probes were purchased from Empire Genomics (Empire Genomics LLC, Buffalo, NY, USA). Slide preparation and hybridization were performed according to the manufacturer’s instructions, and cells (separated by trypsinization from the co-culture) were mounted on coverslips using DAPI/Antifade. Images were captured with a Zeiss Axioplan 2 fluorescence microscope equipped with CytoVision software (Genentix Corp, San Jose, USA). The filter ranges used were XX-green-dUTP (491 nm-516 nm) and XX-orange-dUTP (525 nm-551 nm) for human cells, whereas XX-aqua probes (Aqu-dUTP, 418 nm-467 nm) were used for mouse cells.

**Cardiomyogenic differentiation of CDX2mCherry cells via temporal regulation of Wnt signaling in a feeder-free condition**

CDX2mCherry cells (passages 2-3) underwent cardiac differentiation using a modified GiWi method that involves small molecule Wnt inhibitors (Lian, X et al., Nat Protoc, 2013). H9 huES cells were cultured on Geltrex-coated 6-well plates in mTeSR1 medium until they reached 80–90% confluence. The cells were seeded with 2 mL of mTeSR1 plus 50 nM Chroman in a Geltrex-coated 6-well plate and maintained in a 37°C, 5% CO2 incubator. CDX2mCherry cells, at 70% confluence, were included in the differentiation on day 0. On day 0, 7 μM CHIR99021 (a compound that inhibits GSK3β and activates Wnt signaling) in RPMI/B27-insulin was added to each well for mesoderm induction. Twenty-four hours later, the media were replaced with RPMI/B27-insulin. Seventy-two hours after adding CHIR99021, a working stock of ‘combined medium’ was prepared by mixing half the volume of the medium from the plates with an equal volume of fresh RPMI/B27-insulin medium. Then, 5 µM IWP2 (a Wnt inhibitor) was added to the combined medium before plating into each well. On day 5, the medium was aspirated and replaced with RPMI/B27-insulin. From day 7 onward, the medium was aspirated from each well and replaced with RPMI/B27 plus insulin medium every other day. On days 8, 12, and 20 of induction, flow cytometry and immunostaining with the cardiac marker cTnT (Cat#MA5-12960, ThermoFisher) and sarcomeric actinin (Abcam Cat# ab68168) were performed to characterize the differentiated cardiomyocytes (CM) in each group.

**Intracellular Calcium (Ca²⁺) Measurement**

Cytosolic Ca²⁺ levels were measured using the fluorescent calcium indicator Fura-2 AM (#Cat F1221, Life Technologies, Invitrogen). Cells were grown on a 96-well glass-bottom plate and incubated at 37 °C for 30 minutes in Krebs–Ringer Buffer (KRB) containing 1 mM Ca²⁺. The KRB composition included 135 mM NaCl, 5 mM KCl, 1 mM MgSO₄, 0.4 mM K₂HPO₄, 5.5 mM glucose, and 20 mM HEPES, as previously described (Kuchay, S et al., Nature, 2017). The buffer was supplemented with 2.5 µM Fura-2 AM and 0.02% Pluronic Acid F-127 (Sigma-Aldrich). After incubation, the cells were washed and maintained in 1 mM Ca²⁺/KRB. For functional testing, CDX2-differentiated cardiomyocytes were treated with the β-adrenergic agonist isoproterenol (ISO) at a final concentration of 50 µM. Fluorescence measurements were taken using the Varioskan Lux microplate reader, with cells excited alternately at 340 nm and 380 nm. The emitted fluorescence was recorded, and the data are shown as the 340/380 emission ratio.

***Ex vivo* expansion of CDX2mCherry cells in a feeder-free condition**

FACS-isolated CDX2mCherry cells were plated on 2µg/mL laminin-coated (3 hours at 37°C) cell culture plates using IMDM with 10% FBS. The initial seeding density in a 96-well plate ranged from 1 × 10^4 to 2 × 10^4 cells per well. Media were replenished every 4 days until the cells reached confluence. Once confluent, the cells were dissociated with 0.25% trypsin-EDTA (Cat# 25200-256, Gibco, USA) and replated onto a fresh laminin-coated plate. We evaluated the expression of the smooth muscle marker SM22α **(Figures 4d and e)** to assess mesoderm commitment. For this study, only CDX2mCherry cells up to passage 6 are included.

**Bioreactor-based expansion of CDX2mCherry cells**

In a proof-of-concept experiment run, we used the Distek BioOne (Distek Inc., NJ, USA) single-use benchtop bioreactor system (BioOne1250 2L) to carry out expansion of the placental CDX2mCherry cells on microcarriers (Corning Synthamax II) for 10 days. We optimized cell attachment using unselected chorionic cells for 1 day (data not shown) under identical conditions before initiating the CDX2 cell-selected population. 2D monolayer expanded CDX2mCherry cell were mixed with microcarriers (5cm2/mL) preconditioned with culture media (IMDM+10% FBS) making up to 1L in the vessel. An intermittent ‘pulsed’ agitation process was utilized to support the CDX2-microcarrier attachment portion from Hour 0 to Hour 6 (30-minute cycle: 29 minutes static/1 minute at 90 rpm, seeding Density: 7.5 × 10^5 cells/mL at room temperature). 37 °C and 5% C02 were maintained with dissolved oxygen. 200 mL of fresh media was added on day 5, resulting in the final culture volume of 1.2 L. Further optimization, including controlled pH monitoring and a gas sparging/overlay surface aeration strategy, is underway, which can significantly improve cell proliferation. Despite these uncontrolled parameters, the consistent decrease in the dissolved oxygen curve reflected an increase in cell density of CDX2mCherry cells and, hence, higher cellular metabolism.

**Myocardial Infarction and placental CDX2mCherry cell delivery**

MI was induced in 9-week-old male and female NOD/SCID mice by permanently ligating the LAD. Briefly, the mice were anesthetized with isoflurane and perfused with oxygen in the chamber (initially 3% isoflurane, then maintained at 1.5% during surgery) via endotracheal intubation. The chest cavity was opened, and after carefully dissecting the pericardium, the left anterior descending artery (LAD) was ligated using a 9-0 nylon suture. Once ligated, blanching of the lower left ventricle (LV) indicated ischemia. CDX2mCherry cells and mCherry negative cells (100,000 per site) were injected in a 5 µl volume at three evenly spaced sites around the perimeter of the peri-infarct area with a 31G insulin syringe, fitted with an in-house sterile tubing cover that exposed only 1mm of the needle tip, considering the ventricular wall thickness. Vehicle control mice received an equal volume of phosphate-buffered saline in the same manner. The wound was closed with 6-0 sutures, and after removal from intubation, the mice were allowed to recover gradually in a warm environment (Delta Phase Isothermal pads, Braintree Scientific Inc., MA) and monitored closely. Pre-operatively and for three days postoperatively, 0.05 mg/kg of buprenorphine was administered intraperitoneally as an analgesic. Cardiac MRI was performed the following day as a baseline MI, and again one month later. MRI was conducted using a 7T Bruker Biospec 70/30 scanner (Bruker Corporation). MRI analyses were performed by a clinical cardiologist and an MRI specialist who was blinded to the treatment groups and not involved in the study.

**Fibrosis evaluation**

Fibrosis was evaluated using Masson’s Trichrome Stain Kit (Sigma-Aldrich) on 1-month post-MI heart paraffin-embedded tissue sections from NOD/SCID mice. Three transverse sections from each of three mice per group were analyzed. The kit stains differentiate cytoplasm and muscle fibers in red, collagen in blue, and nuclei in black. Briefly, cryosectioned tissues were deparaffinized, hydrated in deionized water, and fixed in Bouin’s fluid at 56°C for one hour. Subsequent staining steps were performed according to the manufacturer’s protocol. Images were captured with a NanoZoomer S210 digital slide scanner at ×20 magnification and viewed using NanoZoomer Digital Pathology Image (NDPI) software. For analysis, regions of interest—such as infarcted, border, and remote zones from Vehicle, mCherry-negative cell group, and CDX2mCherry cell group, as well as similar regions in sham hearts—were selected. All regions were evaluated using Halo Indica Lab software (version 3.6.4134.137 and Halo AI 3.6.4134). High-resolution images were imported into the software, where sections were annotated and specific staining parameters set to segment and measure the areas of interest accurately. The software distinguished red-stained muscle fibers from blue-stained collagen, enabling the calculation of total tissue area and the percentage of red and blue regions.

**Statistical Analysis**

The number of biological and technical replicates is indicated in the figure legends where appropriate. Data are shown as mean ± SEM unless specified otherwise. ImageJ was used to quantify all immunohistochemistry micrographs and gel images. Matrigel H&E images were quantified using NanoZoomer S210 Digital Slide Scanner at ×20 magnification. Data involving more than two groups were analyzed with one-way ANOVA. For pairwise comparisons, significance was assessed, followed by post-hoc Tukey's or Bonferroni's test to control for inflation of the Type I error rate. An unpaired t-test was applied to analyze the remaining datasets. A two-tailed alpha level of 0.05 was used, and a P value of ≤0.05 was considered statistically significant.

**Data availability**

No unique reagents were generated from this study. Transcriptomics data is deposited at recommended repositories like NCBI GEO, and Mendely (GSE300938, DOI:10.17632/csmzx3crp6.1, DOI: 10.17632/j5v4g89drx.1), and the identifiers are provided in the key resource table. De-identified placental information, along with the patient’s age and race/ethnicity data, will be available upon request. This paper also analyzes existing, publicly available data, accessible at GSE116555 and GSM5537050. All available data reported in this paper will be shared by the lead contact upon request.
