## Supplemental figures and legends for "A multipotent cell type from term human placenta"

†Co-corresponding authors:

**Extended Data Fig.1**

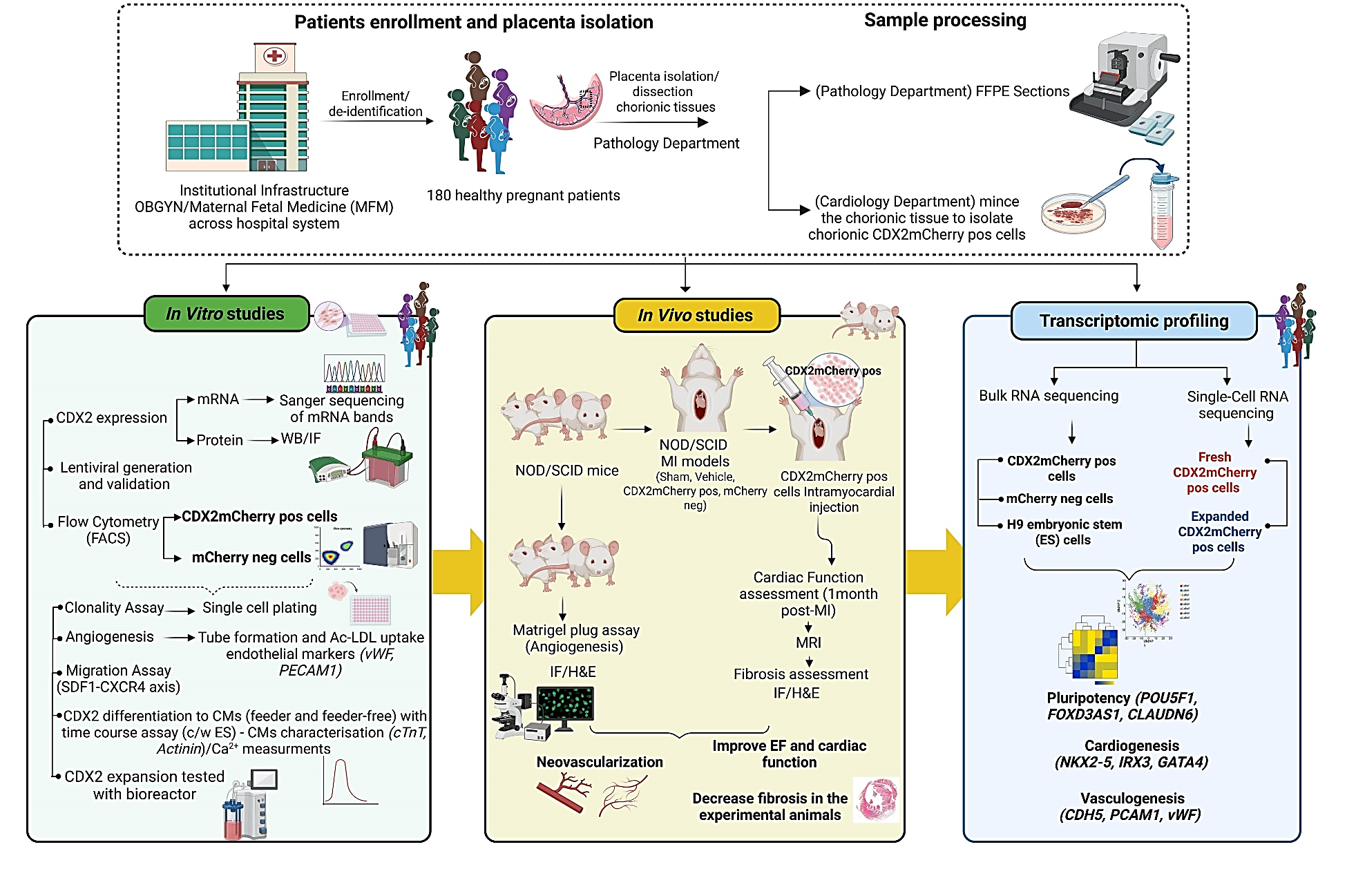

**Extended Data Fig.1:** Schematic overview of the experimental approach used to assess the cardiovascular regenerative potential of placental CDX2 cells

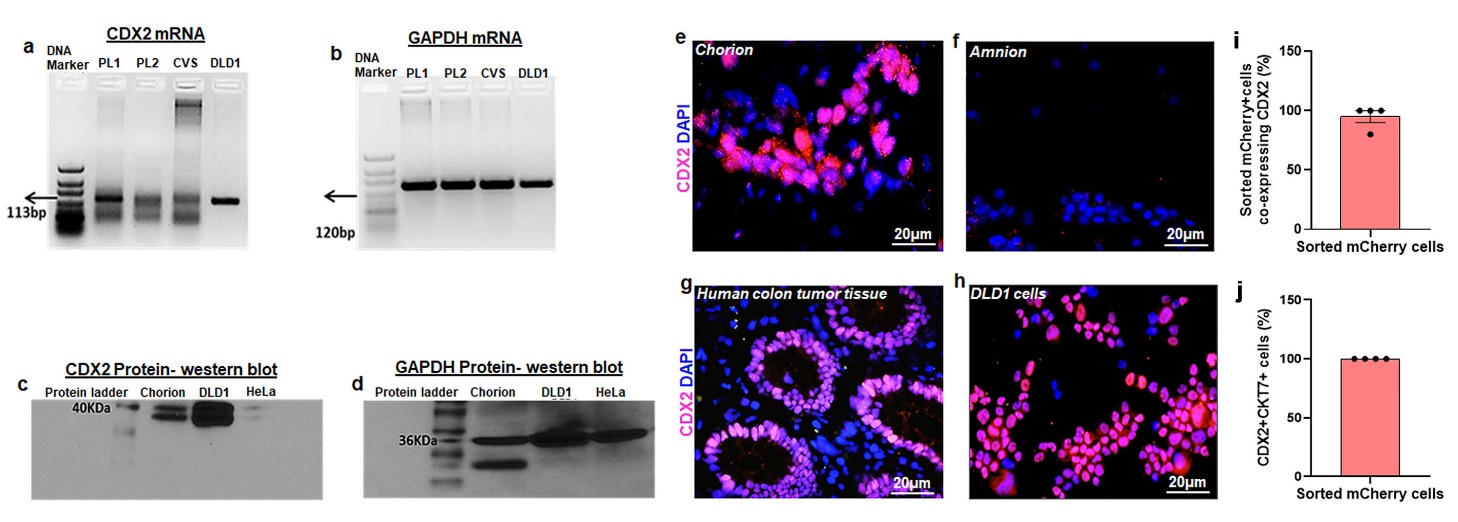

**Extended Data Fig.2**

**Extended Data Fig.2: CDX2 expression in the chorionic region in the human term placenta.**

**a, b)** mRNA raw data related to Figure1a. **c, d)** Western blot raw data of CDX2 protein and GAPDH related to Figure 1b. **e)** Immunofluorescence images of chorionic tissue sections showing nuclear CDX2 expression (pink), nuclei are counterstained using DAPI (blue). **f)** Immunofluorescence images of amnion tissue section showing lack of CDX2 expression. Images represent immunostaining from two different placenta samples. **g)** Immunofluorescence images of human colon tumor tissue. **h)** Human adenocarcinoma cell line DLD1 showed a higher endogenous CDX2 expression and served as a positive control. **i)** Bar plot showing quantification of sorted mCherry cells co-expressing CDX2, detected by immunostaining with anti-CDX2 antibody; related to Figure 2g. **j)** Bar plot showing quantification of sorted mCherry cells that co-express pan trophoblast marker Cytokeratin 7 (CKT7). Data are represented as Mean ± SEM, n=4.

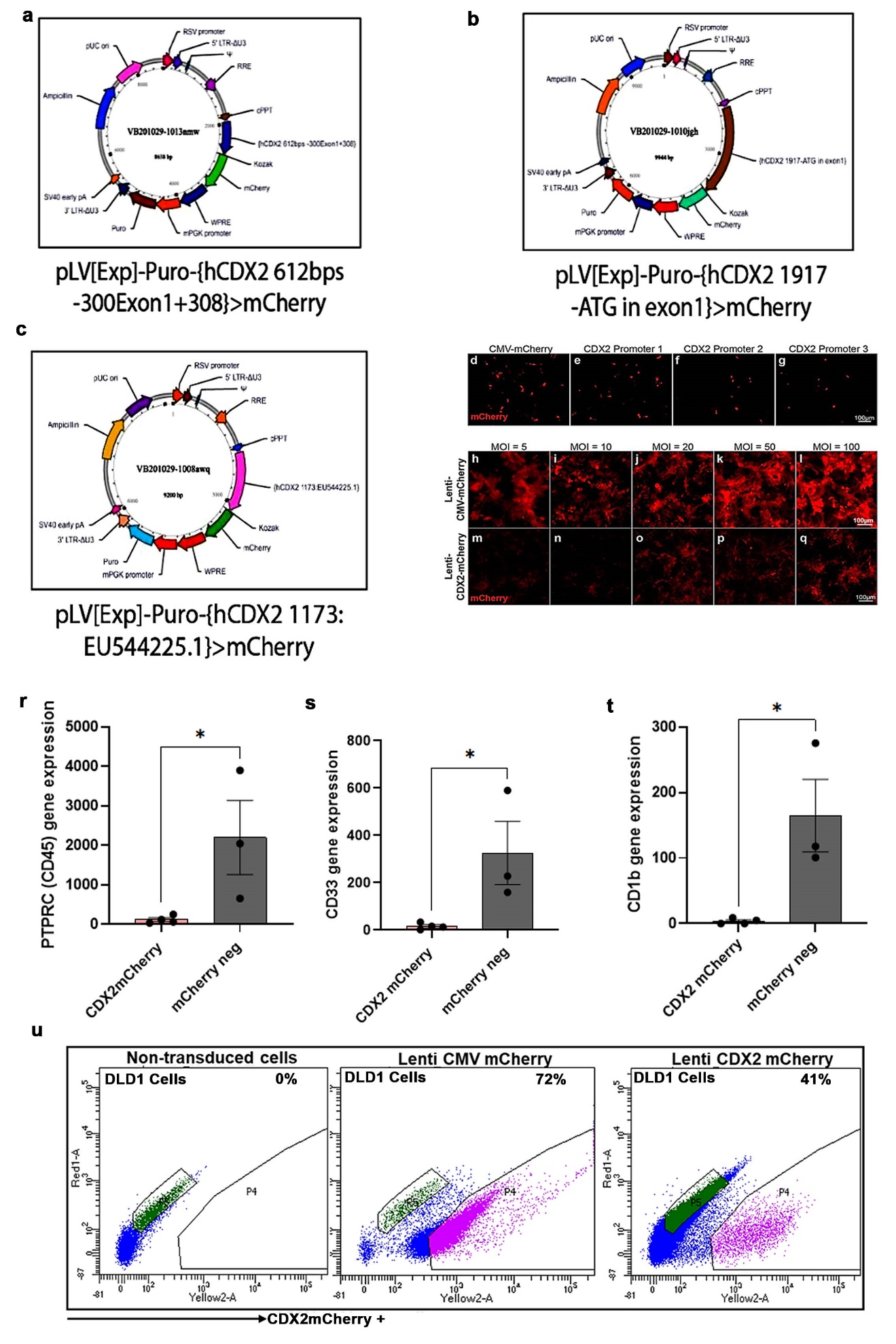

**Extended Data Fig.3**

**Extended Data Fig.3: Human CDX2 promoter-driven lentivirus based isolation and validation in CDX2-expressing and non-expressing cells.**

**a-c)** Vector map incorporating different sequence lengths of huCDX2 promoters. Chosen promoter 1 (CDX2 612bps), is designed to incorporate ~300 sequences upstream of exon 1 and further ~308 bps into exon 1 right before the first open reading frame (ORF) at the ATG (start codon) site. All vector maps containing mCherry were inserted at the ORF as a fluorescence-based readout that reflects the activity of the promoter. **d d-g)** Human adenocarcinoma cell line DLD1 that expresses a higher endogenous CDX2 was used to examine the efficiency of the designed promoters by visualizing the mCherry fluorescence. The plasmid vector showing Promoter 1 **e)** showed an optimal mCherry readout. **h- i)** Lenti CMV mCherry showed an increase in mCherry fluorescence as multiplicity of infection (MOI) was increased from 5-100. **m- q)** CDX2 lenti mCherry (promoter 1) with MOIs ranging from 5-50MOI. **r, s, t)** Bar plots showing flow cytometry-based quantification of mCherry negative cells favoring a hematopoietic lineage with significantly higher expression of CD45, CD33, and antigen-presenting CD1b compared to CDX2 cells. Data are presented as mean ± SEM, CDX2mCherry n=4, mCherry negative n=3 *p≤ 0.05. **u)** Transduction efficiency (50MOI) of selected lenti CDX2 (promoter1) mCherry analyzed by flow cytometry. Left panel: non-transduced cells, middle panel: lenti CMVmCherry, and right panel: lenti CDX2mCherry in DLD1 cells.

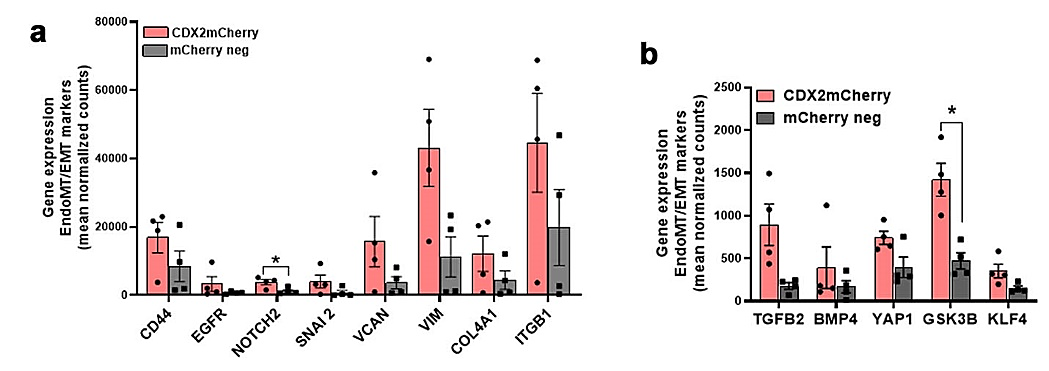

**Extended Data Fig. 4**

**Extended Data Fig.4: Upregulated EndoMT/EMT transcriptome and enriched pathways in CDX2 cells from bulk RNAseq analyses.**

**a, b)** Upregulated markers implicated in the endothelial-mesenchymal transition, a process crucial for valve formation during cardiac development. CDX2 cells showed a significantly higher *NOTCH2* (a) and *GSK3β (b)* gene expression in comparison to mCherry negative cells. Mean normalized values are plotted in the Y-axis, and Log2 fold change in comparison to mCherry negative cells are annotated above each CDX2 bar graph, *p<0.05, n=4.

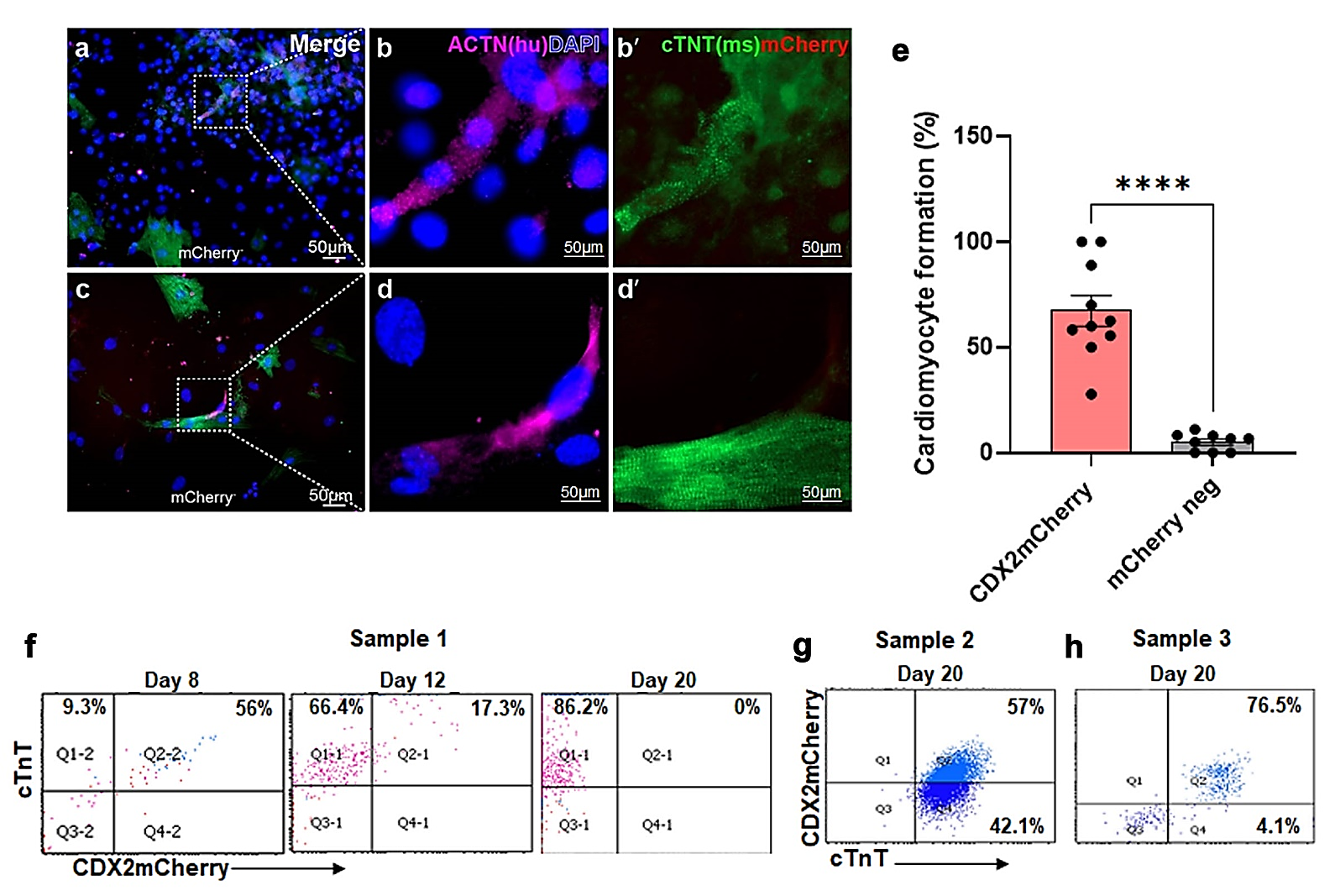

**Extended Data Fig. 5**

**Extended Data Fig.5:** ***In vitro* Cardiac differentiation of CDX2 cells and mCherry negative cells.**

**a, c)** Representative immunofluorescence images from two different mCherry negativeative placenta cells on murine neonatal cardiomyocyte feeders. **b)** Highlighted field from panel a), showing human-specific actinin (pink) and **b’)** cTnT (green) expression. **d)** Highlighted field from panel c) overlay image. **d’)** murine cardiomyocyte feeder with cTnT expression. **e)** Bar plots showing quantification of cardiomyocytes formation wherein CDX2mCherry cells showed significantly higher cardiomyocytes compared to mCherry negative cells. Data are represented as mean ± SEM. CDX2mCherry, n = 5 (2 ROIs per sample), mCherry negativeative, n = 3 (3 ROIs per sample); ****p < 0.0001. **f)** Representative flow cytometry profiles showing time course of differentiation by CDX2mCherry cells (feeder-free method) into cTnT expressing cells downregulating mCherry fluorescence from day 8 to day 20. **g, h)** cytometry profile from two different CDX2 cell populations still retaining mCherry expression (mCherry+cTnT+) on day20 of differentiation indicating inter-sample variability among placental samples.

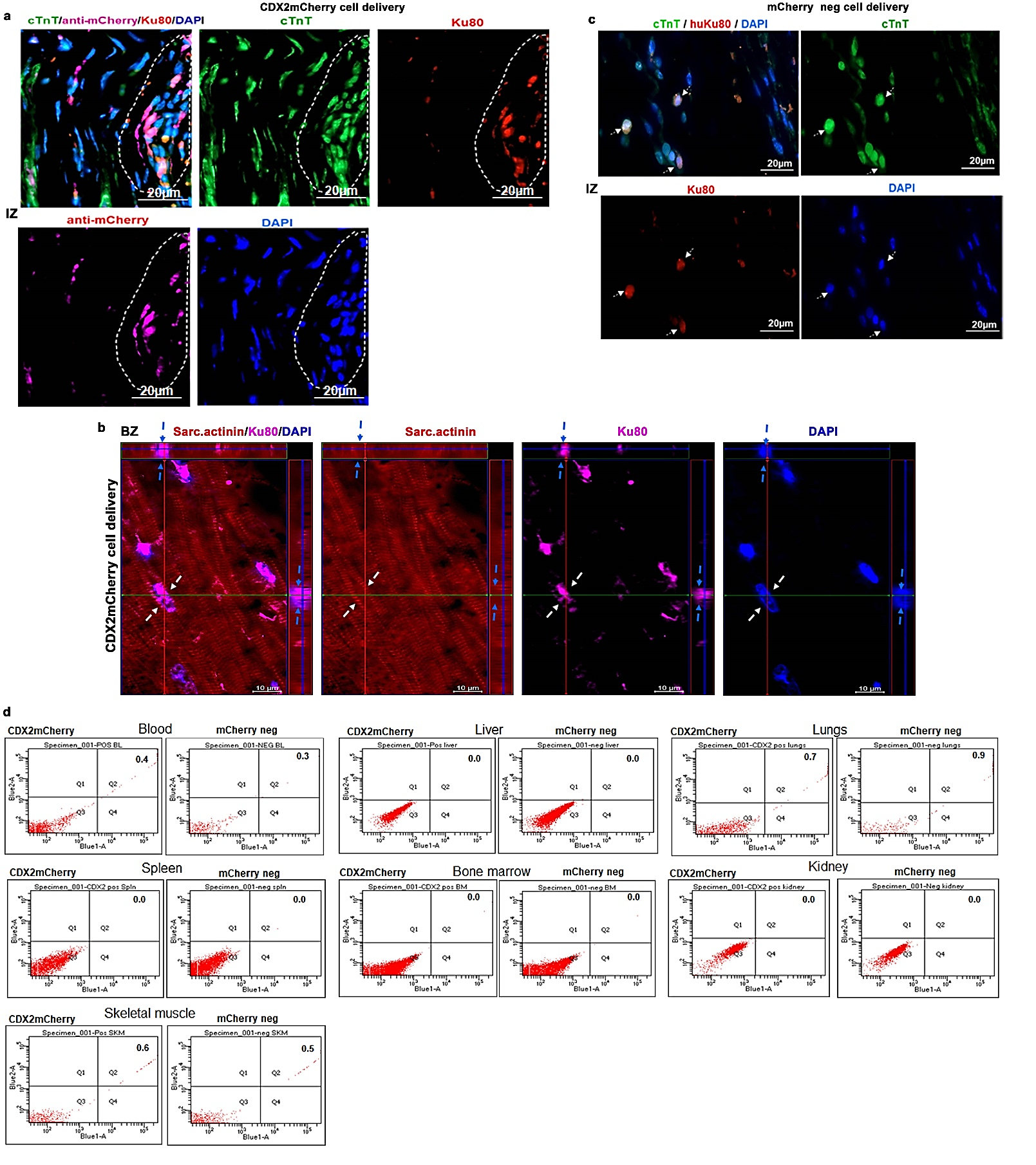

**Extended Data Fig.6**

**Extended Data Fig.6: CDX2 cells differentiated into cardiomyocytes in the infarcted heart.**

**a)** CDX2 cells (dotted lines) in injured myocardium (infarct zone) immunostained for anti-mCherry (pink) and human Ku80 (red), showing cardiac troponin T (cTnT- green). **b)** 3D orthogonal Z-stack of CDX2-derived cardiomyocytes stained for sarcomeric actinin (red) and huKu80 (pink). White arrows indicate sarcomeres. DAPI-stained nuclei (blue) and Ku80+ nuclei (pink) are shown in the Y and Z planes. Nuclei are counterstained with DAPI (blue), related to Figure 6l. IZ= Infarct Zone, BZ= Border Zone. **c)** mCherry negative cells (detected against Ku80-red, and cTnT (green) in the infarct zone. **d)** Representative flow cytometry analyses of cells collected from various non-injured organs of the mice receiving either CDX2 (pos) or mCherry negative cells (neg) showed absence of Ku80+ (quadrant 4, Q4) or mCherry+Ku80+ cells (quadrant 2, Q2).

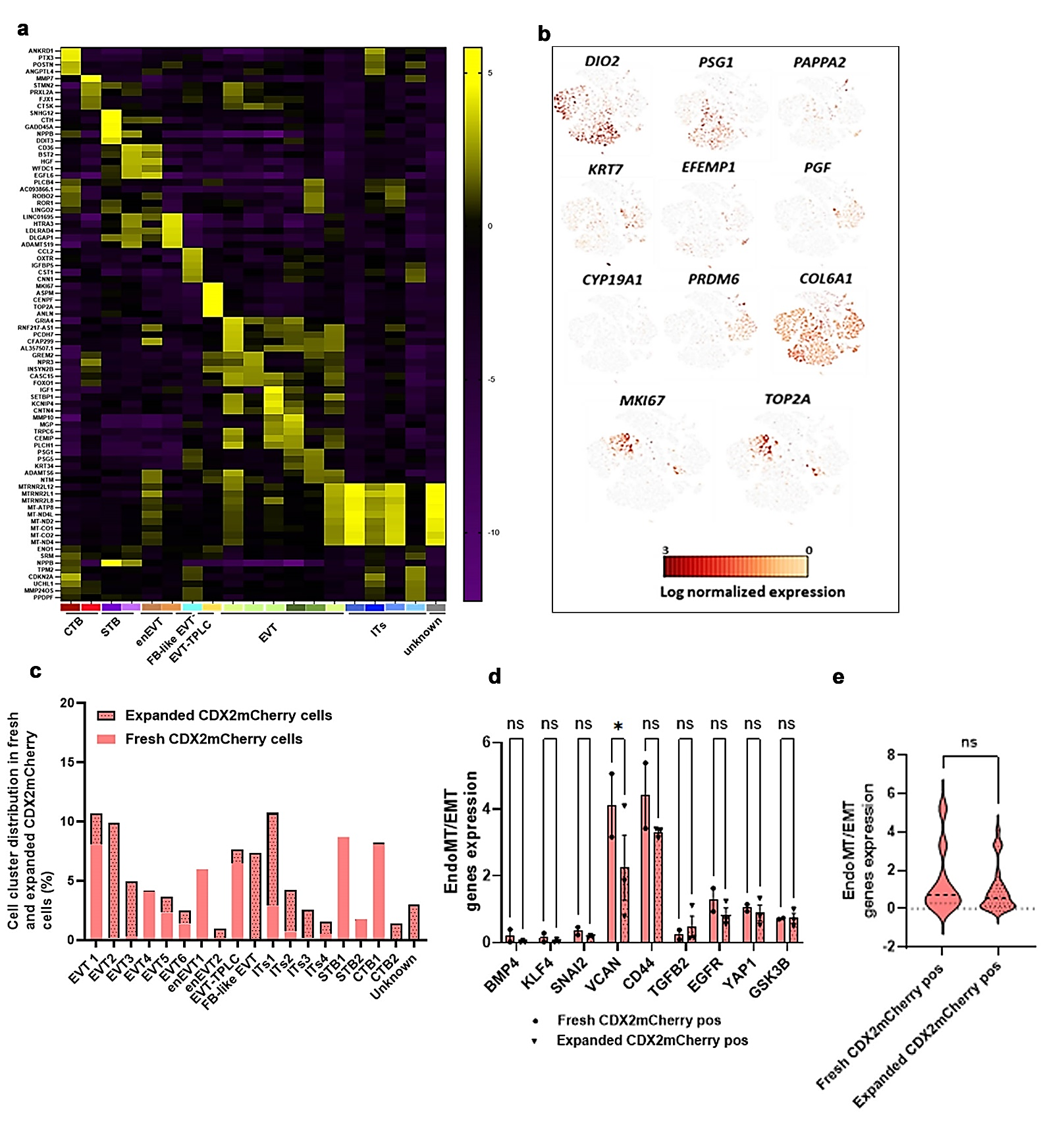

**Extended Data Fig. 7**

**Extended Data Fig.7: scRNA-seq and cluster identification of CDX2 cells.**

**a)** Heatmap displaying the expression levels of the top upregulated genes in the different clusters of the CDX2 cells facilitating the demarcation of the clusters. **b)** t-SNE demonstrating the selected specific gene markers of the EVT (*PSG1, DIO2*), enEVT (*PRDM6*), FB-like EVT (*COL1A1*), EVT-TPLC (*MKI67, TOP2A*), CTB (*KRT7, EFEMP1*), and STB (*PGF*), that were utilized. **c)** Percentage of the cluster-based cell count in fresh and expanded CDX2 cells. **d)** endoMT/EMT gene markers in both fresh and expanded human CDX2 cells from scRNA-seq data corroborating the bulk RNAseq, and **e)** Cumulative expression of endoMT/EMT gene markers in CDX2 cells, showing retention of these markers after expansion. Each dot represents a gene expression value across fresh (n = 2) and expanded (n = 3) samples.

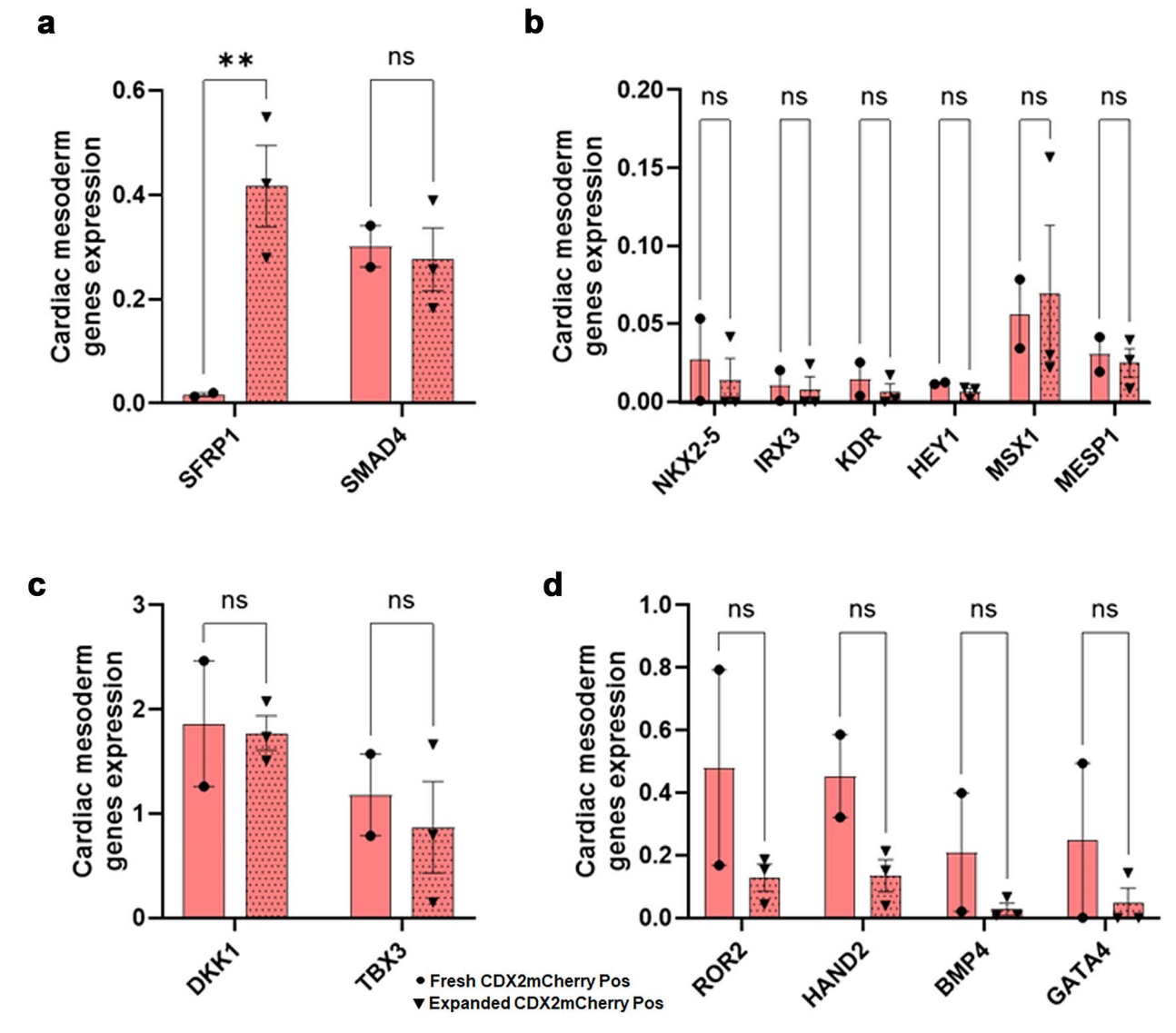

**Extended Data Fig. 8**

**Extended Data Fig.8: Representative bar graphs showing the cardiogenic genes expression.**

**a)** *SFRP1, SMAD4,* and **b)** *NKX2-5, IRX3, KDR, HEY1, MSX1, MESP1,* and **c)** *DKK1, TBX3,* and **d)** *ROR2, HAND2, BMP4, and GATA4* in both CDX2 fresh and expanded CDX2 cells. Except for *SFRP1* (**p<0.01) the other genes did not show any significant change after expansion of CDX2 cells. Data are represented as mean ± SEM (Fresh n=2, expanded n=3 samples).

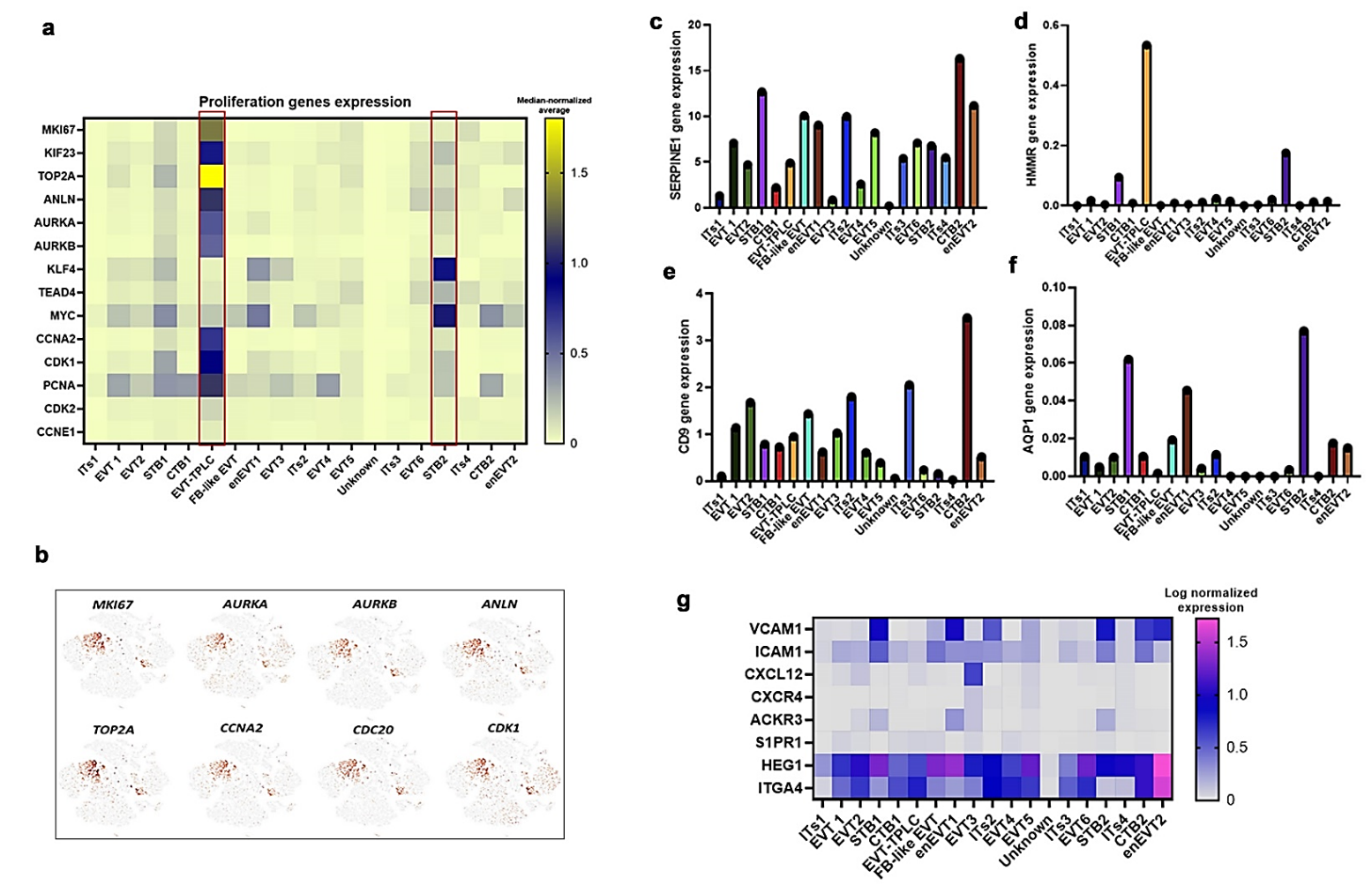

**Extended Data Fig.9**

**Extended Data Fig.9: scRNA-seq identified unique clusters of cells expressing proliferative and homing genes within CDX2 cells.**

**a)** Heatmap representing the expression levels of proliferation genes in different clusters of the CDX2 cells. The two clusters, EVT-TPLC and STB2, are prominent. **b)** t-SNE view of the selected proliferation gene markers (*MKI67, TOP2A, AURKA, AURKB, ANLN, CCNA2, CDC20, CDK1*). **c)** Bar graph representing the surface gene markers *SERPINE,* **d)** *HMMR,* **e)** *CD9* and **f)** *AQP1,* in the different clusters of CDX2 cells. **g)** Heatmap showing the homing gene markers (*ICAM1, CXCL12, CXCR4, ACKR3, ITGA4, HEG1, VCAM1*) in the identified clusters. n=5.

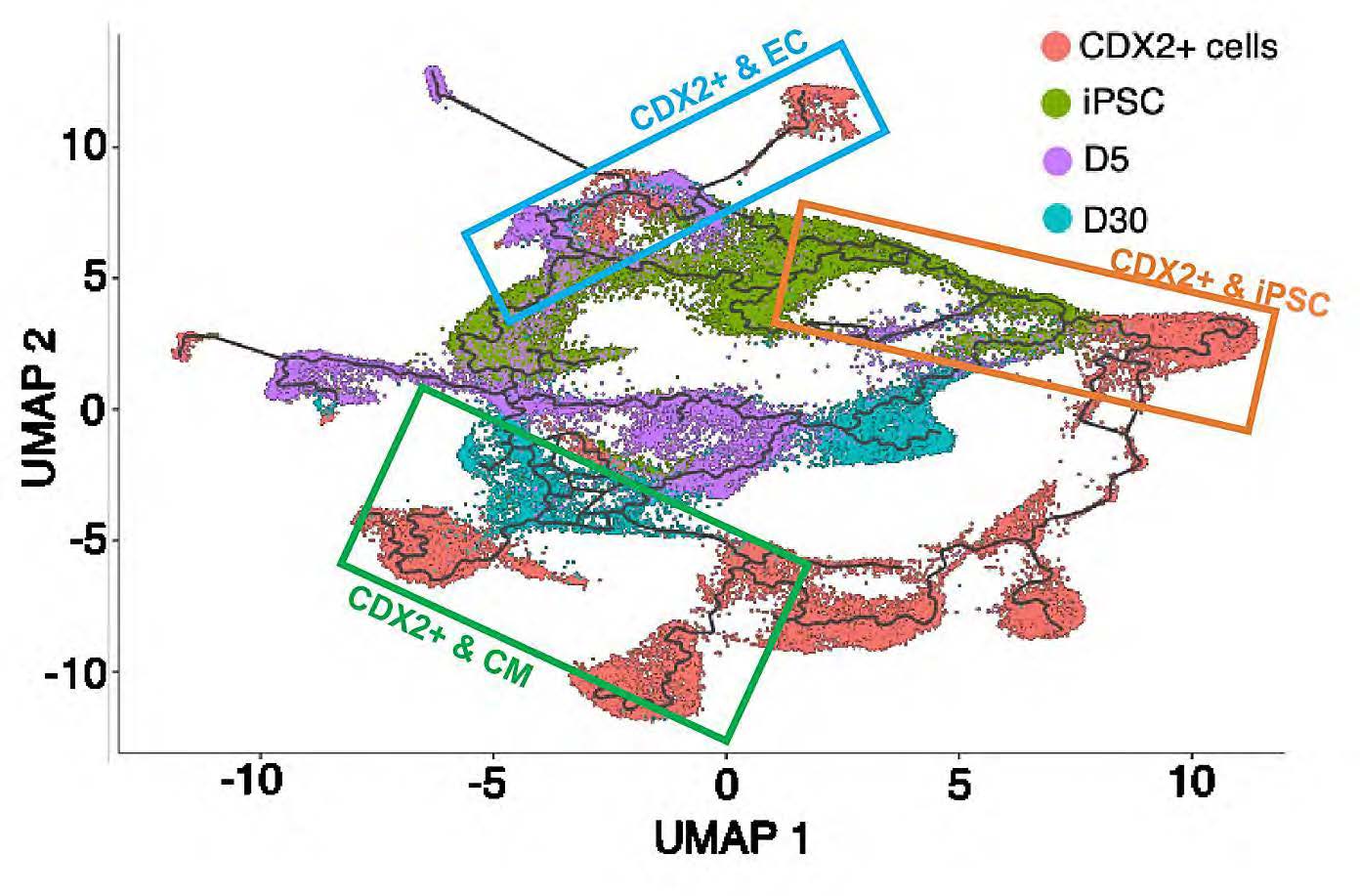

**Extended Data Fig. 10**

**Extended Data Fig.10: Pseudotime trajectory integration of CDX2 cells with iPSC pathways for cardiomyocytes and endothelial differentiation.**

Pseudotime trajectory reveals progression from iPSCs to more differentiated cardiomyocytes, with CDX2 cells mapping along this trajectory. CDX2 cells located near the start of the pseudotime trajectory alongside the iPSCs (Orange box), while cells further along the trajectory were transcriptionally closer to endothelial cells at D5 (Blue box) or cardiomyocytes at D30 (Green box). These findings suggest that CDX2 cells exhibit transcriptional trajectories consistent with progression toward endothelial and cardiomyocyte fates, exhibiting heterogeneity in the differentiation potential.
