## Supplemental tables and guide for "A multipotent cell type from term human placenta"

†Co-corresponding authors:

**Supplementary Table 1: Human CDX2 primer sequences**

| Human CDX2 Sanger sequencing Primer Sequence | Forward GCCACCATGTACGTGAGCTA |
| --- | --- |
| Human CDX2 primers RT PCR | Forward AGGACGTGAGCATGTACCCT  Reverse CACGTGGTAACCGCCGTAG |

**Supplementary Table 2. Summary of differentially expressed genes (DEGs)**

| **Comparison** | **Upregulated genes** | **Downregulated genes** | **Total DEGs** |
| --- | --- | --- | --- |
| **CDX2mCherry+ vs. ESC** | 154 | 1,065 | 1,219 |
| **CDX2mCherry+ vs. mCherry neg** | 529 | 46 | 575 |

**Supplementary Table 3: Left ventricular Ejection Fraction (LVEF) of NOD/SCID mice at baseline post-MI and 1 mo post-MI assessed by MRI**

| **CDX2 cohort (mCherry+ cells)** | **Baseline EF (%) - day1 post MI and injection** | **1-month post-MI EF (%)** |
| --- | --- | --- |
| Mouse1 | 27.9 | 49.7 |
| Mouse 2 | 48 | 51.74 |
| Mouse 3 | 41 | 48.1 |
| Mouse 4 | 47.3 | 53.1 |
| Mouse 5 | 41 | 43.7 |
| Mouse 6 | 38 | 39.02 |
| Mouse 7 | 36 | 45.5 |
| **mCherry neg cohort (mCherry neg cells)** | **Baseline EF (%) - day1 post MI and injection** | **1-month post-MI EF (%)** |
| Mouse1 | 30.1 | 20.8 |
| Mouse 2 | 38.3 | 36.0 |
| Mouse 3 | 45.6 | 29.05 |
| Mouse 4 | 34.3 | 46.46 |
| Mouse 5 | 46.1 | 27.42 |
| Mouse 6 | 38 | 31 |
| Mouse 7 | 32.9 | 22.4 |
| **Vehicle control (PBS)** | **Baseline EF (%) - day1 post MI and injection** | **1-month post-MI EF (%)** |
| Mouse1 | 39.6 | 38.6 |
| Mouse 2 | 38.6 | 23.16 |
| Mouse 3 | 45.5 | 31 |
| Mouse 4 | 27.4 | 15.8 |
| Mouse 5 | 34.1 | 30.6 |
| Mouse 6 | 31 | 27.9 |
| **Healthy mice** | **EF (%)** |  |
| Mouse1 | 64 |  |
| Mouse 2 | 57 |  |
| Mouse 3 | 58 |  |

**Additional Supplemental Data Index**

**Movie S1:** Human placental CDX2mCherry cells forming an endothelial network on Matrigel tube formation assay

**Movie S2:** Endothelial tube formation by HUVEC (positive control) under identical conditions.

**Movie S3:** Day 5 of co-culture demonstrating acquisition of spontaneous beating by CDX2mCherry organoid-like cluster plated on neonatal murine cardiomyocytes feeder.

**Movie S4:** Day 21 of co-culture demonstrating beating of adhered and rod-shaped CDX2mCherry cells on a neonatal murine cardiomyocyte feeder.

**Movie S5:** Cardiac MRI representing the long axis view of CDX2mCherry cell-treated mouse heart at day 1 after MI and cell injection (baseline).

**Movie S6:** Cardiac MRI representing the long axis view of the CDX2mCherry cell-treated mouse heart at 1-month post-treatment.

**Movie S7:** Cardiac MRI representing the long axis view of mCherry negative cell-treated mouse heart at day 1 after MI and cell injection (baseline).

**Movie S8**: Cardiac MRI representing the long axis view of the mCherry negative cell-treated mouse heart at 1-month post-treatment.

**Movie S9:** Cardiac MRI representing the long axis view of the vehicle (PBS injection) treated mouse heart at day 1 after MI and cell injection (baseline).

**Movie S10**: Cardiac MRI representing the long axis view of the vehicle (PBS injection) treated mouse heart at 1-month post-treatment.

**Movie S11:** Cardiac MRI representing the long axis view of the sham mouse heart (no MI) on day 1.

**Movie S12:** Cardiac MRI representing the long-axis view of the sham mouse heart (no MI) at 1 month.

**Source Data Fig.1 (Separate file)**

Sanger sequencing PEAKON data of human CDX2, Sanger sequencing FASTA sequence, corroborating with human CDX2 mRNA related to Fig. 1a

**Source Data Fig. 3a. (Separate Excel file)**

RNAseq raw data Excel spreadsheet showing differentially expressed genes in CDX2mCherry positive cells versus H9 ES cells related to Fig. 3

**Source Data Fig. 3b (Separate Excel file)**

RNAseq raw data Excel spreadsheet showing differentially expressed genes in CDX2mCherry positive cells versus mCherry negative cells related to Fig. 3

**Source Data Fig. 6a (Separate Excel file)**

scRNA-seq raw data Excel spreadsheet showing differentially expressed genes within 19 clusters of the CDX2 cell population related to Fig.6

**Source Data Fig. 6b (Separate Excel file)**

scRNAseq data for cardiogenic, angiovascular, and immune modulatory gene lists related to Fig. 6m,o, and p panels.

**Source Data Fig. 6c** **(Separate Excel file)**

scRNA-seq data Excel file of transcript lists for CDX2 cells, iPSCs, and H9 ES cells related to Fig. 6q.
